# Kv12-Encoded K^+^ Channels Drive the Day-Night Switch in the Repetitive Firing Rates of SCN Neurons

**DOI:** 10.1101/2023.01.30.526323

**Authors:** Tracey O. Hermanstyne, Nien-Du Yang, Daniel Granados-Fuentes, Xiaofan Li, Rebecca L. Mellor, Timothy Jegla, Erik D. Herzog, Jeanne M. Nerbonne

## Abstract

Considerable evidence suggests that day-night rhythms in the functional expression of subthreshold potassium (K^+^) channels regulate daily oscillations in the rates of spontaneous action potential firing of neurons in the suprachiasmatic nucleus (SCN), the master circadian pacemaker in mammals. The K^+^ conductance(s) driving these daily rhythms in repetitive firing rates, however, have not been identified. To test the hypothesis that subthreshold Kv12.1/Kv12.2-encoded K^+^ channels play a role, we obtained current-clamp recordings from SCN neurons in slices prepared from adult mice harboring targeted disruptions in the *Kcnh8* (Kv12.1^−/−^) or *Kcnh3* (Kv12.2^−/−^) locus. We found that mean nighttime repetitive firing rates were higher in Kv12.1^−/−^ and Kv12.2^−/−^, than in wild type (WT), SCN neurons. In marked contrast, mean daytime repetitive firing rates were similar in Kv12.1^−/−^, Kv12.2^−/−^ and WT SCN neurons, and the day-night difference in mean repetitive firing rates, a hallmark feature of WT SCN neurons, was eliminated in Kv12.1^−/−^ and Kv12.2^−/−^ SCN neurons. Similar results were obtained with *in vivo* shRNA-mediated acute knockdown of Kv12.1 or Kv12.2 in adult SCN neurons. Voltage-clamp experiments revealed that Kv12-encoded current densities in WT SCN neurons are higher at night than during the day. In addition, pharmacological block of Kv12-encoded currents increased the mean repetitive firing rate of nighttime, but not daytime, in WT SCN neurons. Dynamic clamp-mediated subtraction of modeled Kv12-encoded currents also selectively increased the mean repetitive firing rates of nighttime WT SCN neurons. Despite the elimination of nighttime decrease in the mean repetitive firing rates of SCN neurons, however, locomotor (wheel-running) activity remained rhythmic in Kv12.1^−/−^, Kv12.2^−/−^, Kv12.1-targeted shRNA-expressing, and Kv12.2-targeted shRNA-expressing animals.

## Introduction

Neurons in the suprachiasmatic nucleus (SCN), the master circadian pacemaker in mammals, including humans, display day-night oscillations in spontaneous repetitive firing rates that drive daily rhythms in physiology and behavior.^[1–4]^ *In vivo* recordings from the rat SCN, isolated from synaptic and humoral inputs, were the first to demonstrate cell autonomous circadian rhythms in the electrical activity of the SCN, with neuronal firing rates higher during the day than at night.^[5]^ Numerous subsequent studies have confirmed that the repetitive firing rates of SCN neurons are higher during the day than at night, with daytime firing rates averaging about 5 Hz and nighttime firing rates averaging about 1 Hz.^[6–10]^ These daily rhythms in repetitive firing rates are accompanied by alterations in input resistances (higher during the day than at night) and membrane potentials (more depolarized during the day than at night), linked to daily changes in subthreshold potassium (K^+^) conductance(s).^[6–10]^ An alternative, “bicycle” model for the circadian regulation of membrane excitability in the SCN, involving a daytime increase in an inward Na^+^ leak current (causing membrane depolarization and increased firing) and a nighttime increase in a K^+^ current (causing membrane hyperpolarization and decreased firing), has also been proposed.^[11]^ The daytime depolarization in the membrane potentials of SCN neurons, however, is associated with increased, *not* decreased, input resistance, as would result from opening inward Na^+^ leak channels. Even in the “bicycle” model,^[11]^ therefore, a reduction in daytime K^+^ conductance is required. Interestingly, this daily pattern (higher firing rates and input resistances during the day than at night) is observed in both diurnal *and* nocturnal animals^[12, 13]^, suggesting that the underlying molecular mechanism(s) is conserved.

Although the daily oscillations in the membrane potentials and input resistances of SCN neurons indicate that day-night changes in subthreshold K^+^ conductance(s) are *required* for rhythmic changes in repetitive firing rates, the critical K^+^ conductance(s) has not been identified.^[7–10]^ Several studies have identified roles for voltage-gated K^+^ (Kv) channels encoded by the *Kcnc1* (Kv3.1),^[14, 15]^ *Kcnc2* (Kv3.2),^[14, 15]^ *Kcna4* (Kv1.4), ^[16, 17]^ *Kcnd1* (Kv4.1)^[18, 19]^ and *Kcnd2* (Kv4.2)^[16–18]^ subunits and for large conductance voltage- and Ca^2+^-dependent K^+^ (BK) channels^[20–22]^ in regulating the repetitive firing rates of SCN neurons. Although day-night differences in the densities of several of these K^+^ currents^[14, 15, 18–20]^ and in the expression levels of transcripts (http://circadb.hogeneschlab.org) encoded by the underlying K^+^ channel subunit genes^[14, 20]^ have been reported, none of these K^+^ channels has been shown to control the day-night switch in the input resistances, membrane potentials and spontaneous repetitive firing rates of SCN neurons.^[7–10]^ The K^+^ conductance(s) driving the cell autonomous circadian rhythms in the repetitive firing properties of SCN neurons remain to be identified. Similar to other types of central and peripheral neurons, *in situ* hybridization and quantitative RNA expression profiling data show that transcripts encoding a number of other K^+^ channel α subunits are expressed in the SCN.^[10, 23]^ Of particular note are two members of the *Kcnh* subfamily, *Kcnh8* (Kv12.1) and *Kcnh3* (Kv12.2), which encode the Kv12.1 and Kv12.2 α subunits, respectively, and are expressed predominately in brain.^[24–26]^ Heterologous expression of Kv12.1 or Kv12.2 gives rise to outward K^+^ currents that function at subthreshold membrane potentials.^[26–30]^ Interestingly, it has been reported that targeted disruption of the *Kcnh3* (Kv12.2) locus increased the input resistances and depolarized the resting membrane potentials of CA1 hippocampal pyramidal neurons.^[30]^ In addition, smaller currents were required to evoke action potentials and repetitive firing rates were higher in Kv12.2^−/−^, compared with wild-type (WT), CA1 hippocampal neurons.^[30]^ The experiments here were designed to directly test the hypothesis that subthreshold Kv12.1- and/or Kv12.2-encoded K^+^ channels underlie the subthreshold K+ conductance that reduces spontaneous excitability of mouse neurons at night. Combining *in vivo* molecular genetic strategies to manipulate Kv12.1 or Kv12.2 expression with *in vitro* electrophysiological and pharmacological approaches to assess the functional consequences of these manipulations, the experiments detailed here demonstrate a critical role for Kv12-encoded K^+^ channels in regulating the nighttime repetitive firing rates and in controlling the day-night rhythms in the repetitive firing rates of SCN neurons.

## Materials and Methods

All reagents were obtained from Sigma-Aldrich, unless otherwise noted.

### Animals

All procedures involving animals were approved by the Animal Care and Use Committee of Washington University and were conducted in accordance with the United States National Institutes of Health Guidelines for the Care and Use of Laboratory Animals. Wild type mice were C57BL/6J. The generation of the Kv12.2^−/−^ mouse line, harboring a targeted disruption in the *Kcnh3* locus, also maintained in the C57BL/6J background, has been described previously.^[30]^ A similar strategy (illustrated in Supplemental Figure 1) was used to generate the Kv12.1^−/−^ line, lacking *Kcnh8*. Crossing Kv12.1^−/−^ and Kv12.2^−/−^ mice provided the Kv12.1^−/−^/Kv12.2^−/−^ double knockout (DKO) line. All mice were maintained in the C57BL/6J background in one of the Washington University Danforth or Medical School animal facilities.

### Screening of Kv12.1- and Kv12.2-targeted shRNAs

An interfering RNA strategy^[31]^ was developed to allow the acute *in vivo* knockdown of Kv12.1 or Kv12.2 expression selectively in the SCN of adult animals. In initial screening experiments, short hairpin RNA (shRNA) sequences targeting Kv12.1 (n = 5) or Kv12.2 (n = 5), obtained from the RNAi Consortium (TRC) through the Genome Institute at Washington University Medical School, were evaluated *in vitro* to determine the efficiency of Kv12.1 or Kv12.2 knockdown. For screening, tsA201 cells were co-transfected, using PepMute (Signagen), with a cDNA construct encoding Kv12.1-eYFP or Kv12.2-eYFP and one of the Kv12.1-targeted or Kv12.2-targeted shRNAs. Additional experiments were conducted using a non-targeted (NT), control shRNA generated against a variant of green fluorescent protein. Approximately 48 hrs following the transfections, cell lysates were prepared, fractionated by SDS-PAGE, transferred to polyvinylidene fluoride (PVDF) membranes and probed for eYFP expression (Millipore; polyclonal anti-GFP antibody, 1:1000). Blots were also probed with an alpha-tubulin antibody (Abcam; monoclonal anti-α-tubulin, 1:10,000) to verify equal protein loading of each lane. The efficiency of the knockdown of Kv12.1-eYFP or Kv12.2-eYFP by each of the Kv12.1- or Kv12.2-targeted shRNAs was quantified by densitometry. The Kv12.1- and Kv12.2-targeted shRNA sequences producing the largest reduction in Kv12.1-eYFP (5’-CAACATTAACTCAGGAGGTTT-3’) or Kv12.2-eYFP (5’- CAGCTTTATGGACCTCCAC TT-3’) expression were subsequently evaluated for specificity. In these experiments, tsA201 cells were co-transfected with cDNA constructs encoding Kv12.1- eYFP, Kv12.2-eYFP or Kv4.1-eYFP, together with the Kv12.1-targeted shRNA, Kv12.2-targeted shRNA or NT shRNA. Two days after the transfections, lysates were prepared and fractionated. Western blots were probed with the anti-GFP and anti-tubulin antibodies and quantified by densitometry.

### Generation of shRNA-expressing adeno-associated viruses

The selected Kv12.1-targeted, Kv12.2-targeted, and NT shRNA 21-nucleotide sense sequences were synthesized (Integrated DNA Technologies) into the corresponding 97-nucleotide microRNA-adapted shRNA oligonucleotides, containing sense and antisense sequences linked by a 19-nucleotide hairpin loop. Forward and reverse strands were annealed and cloned, in a microRNA (human miR30) context, into the 3’-untranslated region (3’-UTR) of eGFP in the pPRIME vector.^[32]^ The entire eGFP-shRNA cassette was then cloned into a viral shuttle vector with a synapsin (SYN) promoter. Adeno-associated viruses, serotype 8 (AAV8), shown previously to produce robust transduction of adult mouse SCN neurons *in vivo*,^[19]^ were generated by the Hope Center Virus Core Facility at Washington University Medical School.

### Stereotaxic virus injections

Bilateral stereotaxic injections of the NT, the Kv12.1-targeted, or the Kv12.2-targeted shRNA-expressing AAV8 were made into the SCN of adult (6-10 week) wild type (WT) C57BL/6JN male and female mice using previously described methods.^[19]^ Briefly, under sterile conditions, each mouse was anesthetized with isoflurane (3%) and secured in a stereotaxic head frame (Kopf Instruments). The head was shaved and Betadine was applied to cleanse and sterilize the shaved region. An incision was then made along the midline and the skin was pulled back to expose the skull. One of the shRNA-expressing viruses was injected (∼ 600 nl) into each hemisphere of the SCN (coordinates: 0.3 mm rostral to bregma, 0.1 mm left and right to midline and 5.6 mm ventral to pial surface). The injection syringe (Hamilton) delivered the virus at a constant rate of 0.1 μl/min using a syringe pump (KD Scientific). The syringe was left in place for ∼5 min after the injection was completed to minimize the upward reflux of solution during the removal of the needle. Vetabond tissue adhesive (3M, Maplewood, MN) was used to close the incision. Immediately following the surgery, animals were allowed to recover from the anesthesia on a heating pad maintained at 37°C and were given an intraperitoneal (IP) injection of Rimadyl (0.1 ml of 0.05 mg/ml, Pfizer).

### Preparation of SCN slices

SCN slices (300 μm) were prepared from adult (8-12 week old) WT, Kv12.1^−/−^, Kv12.2^−/−^, and Kv12.1^−/−^/Kv12.2^−/−^ (DKO) mice, maintained in either a standard (lights on at 7:00 am and lights off at 7:00 pm) or a reversed (lights on at 7:00 pm and lights off at 7:00 am) 12 hr:12 hr light-dark (LD) cycle, using previously described methods.^[16, 17, 19]^ Zeitgeber times (ZT) are indicated: ZT0 corresponds to the time of lights on and ZT12 to the time of lights off in the animal facility. Daytime slices were routinely prepared at ZT5 from WT, Kv12.1^−/−^, Kv12.2^−/−^ and DKO mice maintained in the standard LD cycle and nighttime slices were routinely prepared at ZT16 from WT, Kv12.1^−/−^, Kv12.2^−/−^ and DKO mice maintained in the reversed LD cycle. Using a similar strategy, SCN slices were also prepared from mice two weeks following NT, Kv12.1-targeted or Kv12.2-targeted shRNA-expressing AAV8 injections into the SCN. In an additional set of experiments designed to determine if firing rates were affected at any time during the cycle, slices were also obtained from Kv12.1^−/−^ and Kv12.2^−/−^ animals during the transition from ‘lights-on to lights-off’ (ZT12-ZT14), as well as slices prepared mid-day (ZT6-ZT8) and mid-evening (ZT18-ZT20).

For the preparation of daytime slices, brains were rapidly removed (in the light) from animals anesthetized with 1.25% Avertin (Acros Organics, 2,2,2-tribromoethanol and tert-amyl alcohol in 0.9% NaCl; 0.025 ml/g body weight) and placed in ice-cold cutting solution containing (in mM): sucrose, 240; KCl, 2.5; NaH_2_PO_4_, 1.25; NaHCO_3_, 25; CaCl_2_, 0.5; and MgCl_2_ 7, saturated with 95% O_2_/5% CO_2_. In separate experiments, some daytime slices were prepared at ZT3 and ZT11 to obtain electrophysiological recordings earlier in the day (ZT6-ZT8) and during the transition from “lights on” to “lights off” (ZT12 - ZT14). For the preparation of nighttime slices, animals in the reversed LD cycle were removed from their cages at ZT15 under infrared illumination, anesthetized with isoflurane and enucleated using previously described procedure.^[33–35]^ Following an IP injection of Rimadyl (0.1 ml of 0.05 mg/ml), each animal was allowed to recover from the anesthesia (for approximately 1 hr) prior to the preparation of slices. At ZT16, animals were anesthetized with 1.25% Avertin; brains were rapidly removed and placed in ice-cold cutting solution. For all experiments, coronal slices (300 µm) were cut on a Leica VT1000 S vibrating blade microtome (Leica Microsystems Inc.) and incubated in a holding chamber with oxygenated artificial cerebrospinal fluid (ACSF) containing (in mM): NaCl, 125; KCl, 2.5; NaH_2_PO_4_, 1.25; NaHCO_3_, 25; CaCl_2_, 2; MgCl_2_, 1; and dextrose, 25 (∼310 mOsmol l^−1^), saturated with 95% O_2_/5% CO_2_, at room temperature (23-25°C) for at least 1 hr before transfer to the recording chamber.

### Electrophysiological recordings

Whole-cell current-clamp and action potential- (voltage-) clamp recordings were obtained at room temperature (23-25°C) during the day (ZT7-ZT12) or at night (ZT18-ZT24) from SCN neurons in slices prepared (as described above) during the day or at night from WT, Kv12.1^−/−^, Kv12.2^−/−^ or DKO (Kv12.1^−/−^/Kv12.2^−/−^) mice. Current-clamp recordings were also obtained from SCN neurons in slices prepared during the day or at night two weeks following NT, Kv12.1-targeted or Kv12.2-targeted shRNA-expressing AAV8 injections. For recordings, SCN neurons were visually identified in slices using differential interference contrast optics with infrared illumination. Slices were perfused continuously with ACSF containing 20 μM Gabazine (Tocris Bioscience) and saturated with 95% O_2_/5% CO_2_. Whole-cell current-clamp recordings were obtained using pipettes (4-7 MΩ) containing (in mM): 144 K-gluconate, 10 HEPES, 3 MgCl_2_, 4 MgATP, 0.2 EGTA and 0.5 NaGTP (pH 7.3; 300 mOsmol l^-1^). Tip potentials were zeroed before recordings were obtained.

For each cell, a loose patch, cell-attached recording was first obtained and the spontaneous repetitive firing of action potentials was recorded for ∼1 min. Following the formation of a GΩ seal, the whole-cell configuration was established, and whole-cell membrane capacitances and series resistances were compensated. Whole-cell spontaneous firing activity was then recorded for ∼1 min. Access resistances were 15-20 MΩ, and data acquisition was terminated if the access resistance increased by *≥* 20%. Voltage signals were acquired at 100 kHz, filtered at 10 kHz and stored for offline analysis. Data were collected using a Multiclamp 700B patch clamp amplifier (Molecular Devices) interfaced to a Dell personal computer with a Digidata 1332 and the pCLAMP 10 software package (Molecular Devices). Consistent with previous studies,^[14, 15, 17, 20, 33, 36]^ these loose-patch (extracellular) and whole-cell (intracellular) recordings revealed that daytime and nighttime adult mouse SCN neurons display tonic and irregular spontaneous repetitive firing patterns. To facilitate comparison of repetitive firing rates across groups, the average repetitive firing rate of each cell during the initial 1 min of recording was determined. Mean ± SEM repetitive firing rates for each group were calculated and are reported here. Input resistances (R_in_) were determined by measuring the steady-state voltage changes produced by ± 5 pA current injections from a membrane potential of -70 mV. The voltage threshold for action potential generation in each cell was determined as the point during the upstroke (depolarizing phase) of the action potential at which the second derivative of the voltage was zero. Resting membrane potentials (V_r_) were estimated from phase plots of spontaneous action potentials.

For action potential- (voltage-) clamp recordings, CdCl_2_ (0.1 mM) and tetrodotoxin (1 μM) were added to the standard ACSF bath solution described above. Recording pipettes (3-5 MΩ) contained (in mM): 144 K-gluconate, 10 HEPES, 3 MgCl_2_, 4 MgATP, 0.2 EGTA and 0.5 NaGTP (pH 7.3; 300 mOsmol l^-1^). Spontaneous repetitive firing activity, recorded from a nighttime WT SCN neuron, was used as the voltage command. Tip potentials were zeroed before membrane-pipette seals were formed. Following formation of a GΩ seal and establishing the whole-cell configuration, membrane capacitances and series resistances were compensated electronically. Series resistances were in the range of 15-20 MΩ, and were routinely compensated by 70-80%. If the series resistance changed ≥ 20% during a recording, the experiment was stopped and acquired data from that cell were not included in the analyses. Outward K^+^ currents evoked by the action potential voltage command in WT and DKO (Kv12.1^−/−^/Kv12.2^−/−^) SCN neurons were recorded in TTX- and CdCl_2_-containing ACSF bath solution before and after local application of 20 μM CX4 (1-(2-chloro-6-methylphenyl)-3-(1,2-diphenylethyl) thiourea)^[30]^ dissolved in the same (TTX- and CdCl_2_-containing) ACSF bath solution. Off-line subtraction of the outward K^+^ currents recorded after local application of CX4 from the currents recorded prior to CX4 application provided the CX4-sensitive K^+^ currents.

Additional experiments were conducted to determine the voltage dependence of activation of the CX4-sensitive currents in nighttime WT SCN neurons. To determine the voltage-dependence of activation of the CX4-sensitive currents, whole-cell Kv currents, evoked in response to 2 sec depolarizing voltage steps to potentials between -100 and +50 mV (in 10 mV increments) from a holding potential (HP) of -70 mV, were first recorded in standard ACSF solution containing tetraethylammonium (10 mM), 4-aminopyridine (10 mM), CdCl_2_ (0.1 mM) and tetrodotoxin (1 μM). Outward Kv currents, evoked using the same protocol, were recorded again (from the same cell) with CX4 (20 μM) added to the same ACSF solution. Off-line digital subtraction of the Kv currents recorded with and without the CX4 in the bath provided the CX4-sensitive currents (I_CX4_) (Supplemental Figure 2A). In each cell, I_CX4_ conductances at each test potential were calculated and normalized to the maximal conductance (G_max_), determined in the same cell. Mean ± SEM normalized I_CX4_ conductances (G/G_max_) were then plotted as a function of the test potential and fitted using the Boltzmann equation, G/G_max_ = 1 + e^[(Va – Vm)/k]^ (Supplemental Figure 2B), where V_a_ is the voltage of half-maximal activation and *k* is the slope factor.

### Modeling Kv12-encoded (I_Kv12_) and A-type (I_A_) currents and Dynamic Clamp Recordings

A three-state Markov model of *I_Kv12_* channel gating, with two closed states, C_1_ and C_2_, and one open state, O, was developed using MATLAB (MathWorks Inc). We used published kinetic and steady-state activation and deactivation data for heterologously expressed Kv12.1-encoded channels.^[24]^

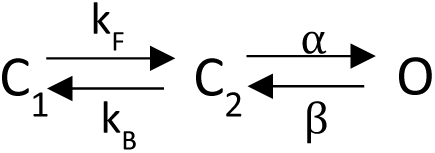

The voltage-dependent forward and reverse transition rates, *α* and *β*, between the C_2_ and O states, determined from the published data,^[24]^ were: *p_1_*exp(*p_2_*V_m_) and *p_3_*exp(*-p_4_*V_m_), respectively, where V_m_ is the membrane voltage, *p*_1_ = 0.019, *p*_2_ = 0.065, *p*_3_ = 0.011, and *p*_4_ = -0.039.^[37]^ The voltage-independent transition rates (*k_F_* and *k_B_)* between the C_1_ and C_2_ states were approximated from the published time constants of slow activation and the ratio of the fast and slow components of activation of Kv12.1-encoded currents.^[24]^ The values determined for *k_F_* and *k_B_* were 0.008 and 0.0013, respectively. The steady-state voltage-dependence of activation of the model was then adjusted using the voltage-clamp data acquired from SCN neurons with a Kv12-selective inhibitor CX4 (see: Supplemental Figures 2A and 2B). The Kv12 current is calculated as: I_Kv12_ = G*P(O)*(V_m_-E_K_) where G is a scalable parameter for current magnitude, P(O) is the probability that the channel is open, and E_K_ is the reversal potential for K^+^.

A Markov model describing the gating of the K^+^ channels that generate the A-current (I_A_) was also developed using MATLAB (MathWorks Inc). The model is based on a previously described model of the rapidly activating and inactivating K^+^ current in (ferret) ventricular myocytes, consists of three closed states C_1_ - C_3_, two inactivated states I_0_ and I_1_, and an open state (O),^[38]^ and was populated using acquired voltage-clamp data detailing the time- and voltage-dependent properties of I_A_ in mouse SCN neurons.^[19]^

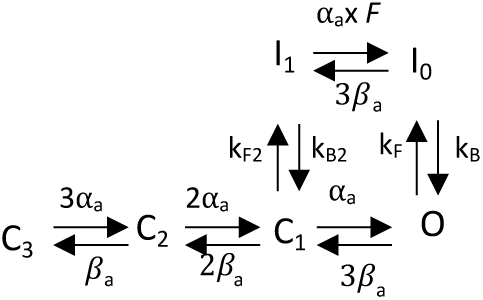

The voltage-dependent transition rates for I_A_ activation and deactivation, *α* and *β*, determined from analyses of the time constants and sigmoidicity of I_A_ activation in SCN neurons^[19]^, were: *p_1_*exp(*p_2_*V_m_) and *p_3_*exp(*-p_2_*V_m_), respectively, where V_m_ is the membrane voltage, *p*_1_ = 0.5, *p*_2_ = 0.0328, and *p*_3_ = 0.0680. In contrast to activation, the time constants of I_A_ inactivation in mouse SCN neurons were voltage independent over a wide range of membrane potentials.^[19]^ This finding suggests a “ball and chain” inactivation mechanism in which a putative ‘inactivation particle’ interacts with the open channel to cause inactivation.^[39]^ In the model, this inactivated state is I_0_, and k_F_, the voltage-insensitive “on” rate for the particle, is equal to the reciprocal of the inactivation time constant at positive potentials. The unbinding rate of the inactivation particle, k_B_, was initially estimated by the ability of the model to match steady-state inactivation data for I_A_.^[40]^ Such a single state “ball and chain” model recovers very slowly from inactivation when the membrane is returned to hyperpolarized potentials. To account for the voltage-insensitive inactivation of I_A_ with relatively rapid recovery from inactivation, as observed experimentally,^[19]^ we introduced a transition state, I_1_, into the model. In contrast to I_0_, the I_1_ state is a low-affinity binding state in which the energy from the backwards movement of the voltage sensor during recovery destabilizes the binding interaction with the inactivation particle in a “push-off ball and chain” inactivation mechanism.^[40]^ The voltage-independent transition rates, *k_F_* (0.05), *k_B_* (0.00025), *k_F2_* (0.05), and *k_B2_* (0.002), were determined by fitting the data describing steady-state inactivation of I_A_, and the experimentally determined time constants of I_A_ inactivation and recovery (at -70 mV) from inactivation.^[19]^ The ratio factor, *F* (8), was determined by macroscopic reversibility. The currents are calculated as: I_A_ = G*P(O)*(V_m_-E_K_), where G is a scalable parameter for current magnitude, P(O) is the probability that the channel is open, and E_K_ is the reversal potential of K^+^.

Dynamic clamp was carried out using a commercially available Cybercyte Dynamic Clamp System, Cybercyte DC1,^[41, 42]^ from Cytocybernetics (Buffalo, NY). This system consists of a 16 channel, 16 bit, 100 kS/s MCC PCIe-DAS1602/16 board installed and configured in a Dell 5820 Precision workstation. The average loop time determined for this system was 22 μs. Prior to experiments on SCN neurons, the I_K12_ and I_A_ formulations were converted to Cybercyte channel definition files using Cybersolver software and validated by applying the simulated currents to a model cell (Molecular Devices) with an input resistance of 500 MΩ and a capacitance of 30 pF in whole-cell mode. The Cybercyte DC1 system allows current amplitudes to be scaled during dynamic clamp experiments such that the modeled currents (I_K12_ and I_A_) can be “added” or “subtracted” (by the addition of I_K12_ or I_A_ of the opposite polarity) in real time during whole-cell current-clamp recordings from daytime or nighttime WT SCN neurons. Preliminary experiments were undertaken to determine the minimal I_Kv12_ (or I_A_) amplitude that, when added/subtracted, changed the spontaneous repetitive firing rates of most (9 of 10) nighttime SCN neurons. These experiments revealed that the minimal subtracted I_Kv12_ amplitude required to affect firing was 2 pA, whereas the minimal added I_Kv12_ amplitude to alter firing was 0.5 pA. For I_A_, the minimal modeled I_A_ current amplitude to alter firing was 20 pA. For dynamic clamp recordings, the membrane voltage was sampled at 20 kHz and the Cybercyte DC1 returned the corresponding modeled I_Kv12_ (or I_A_) at a rate of 50 kHz. For each cell, spontaneous repetitive firing was recorded for 1 min. Modeled currents were then added or subtracted in multiples (i.e., x, 2x, -x, -2x, etc.) of the minimal (x) amplitudes, determined as described above, and the resulting repetitive firing rates were measured. The percent change in the repetitive firing rate with each current injection was then determined in each cell. These values were then averaged across cells and the mean ± SEM percent changes in firing rates as a function of the injected current amplitudes are presented.

### Analyses of wheel-running activity

Adult (9-12 week old) WT, Kv12.1^−/−^, Kv12.2^−/−^ and DKO (Kv12.1^−/−^/Kv12.2^−/−^) mice were placed (individually) in cages equipped with running wheels in light-tight chambers illuminated with fluorescent bulbs (2.4 ± 0.5 × 10^18^ photons/s*m^2^; General Electric). Wheel-running activity was recorded (Clocklab Actimetrics) in 6 min bins for 10 days in a 12:12 hr LD cycle, followed by recordings in constant darkness (DD) for at least 20 days. Similar recordings were obtained from animals 10 days following bilateral injections (600 nl in each hemisphere) of the NT, Kv12.1-targeted or Kv12.2-targeted shRNA-expressing AAV8 into the SCN of adult (9-12 week) WT mice. The period of rhythmicity of each mouse was determined using χ^2^ periodogram analysis^[43]^ of continuous recordings for 10 days in DD (Clocklab). Wheel-running was considered rhythmic if the χ^2^ periodogram value exceeded the 99.9% confidence interval (Qp value). Statistical analysis of the circadian periods were compared between groups using a one-way analysis of variance (ANOVA).

### Quantitative RT-PCR analysis

Total RNA was isolated from (300 μm) SCN slices, collected every 4 hrs (ZT 3, 7, 11, 15, 19 and 23) for 2 consecutive days from adult (8 -10 week) WT mice (n = 7-8), maintained in standard and reversed LD conditions, and DNase treated using previously described methods.^[44]^ RNA concentrations were determined by optical density measurements.^[44]^ The mRNA transcript expression levels of genes encoding *Per2*, *Bmal*, *Kcnh8* (Kv12.1), *Kcnh3* (Kv12.2), as well as of the endogenous control gene *Hprt* (hypoxanthine guanine phosphoribosyl transferase), were determined using Taqman-based real-time quantitative (RT) PCR. Data were collected with instrument spectral compensations using the Applied Biosystems SDS 2.2.2 software and analyzed using the threshold cycle (C_T_) relative quantification method.^[45]^ The expression of each transcript was normalized to the expression of *Hprt* in the same sample and evaluated for rhythmicity using JTK cycle analysis^[46]^ with the period set to 24 hr. The primers sequences used were: *Per2*: 5’-TCCACCGGCTACTGATGCA and 5’- TGGATGATGTCTGGCTCATGA; *Bmal*: 5’- GTAGGATGTGACCGAGGGAAGA and 5’- AGTCAAACAAGCTCTGGCCAAT; *Kcnh8* (Kv12.1): 5’- AGGATTACTGGCGCCACAGA and 5’- CTTTGCCACTTGGGCATTG; *Kcnh3* (Kv12.2): 5’- GCAACGTGTCCGCTAACACA and 5’- GCCGTCACATTCCCAAACA; and *Hprt*: 5’- TGAATCACGTTTGTGTCATTAGTGA and 5’-TTCAACTTGCGCTCATCTTAGG.

### Statistical analysis

Electrophysiological data were compiled and analyzed using ClampFit (v. 10.2, Molecular Devices), Mini Analysis (v. 6.0.7, Synaptosoft, Decatur, GA), and Prism (v. 8.2, GraphPad Software, La Jolla, CA). Averaged data are presented as means ± standard error of the means (SEM). Statistical analyses were performed using the Student’s t-test or one-way analysis of variance (ANOVA) with Newman-Kuels post-hoc pairwise comparisons, as indicated in the text, figure legends, or tables; *P* values are reported.

## Results

### Targeted deletion of Kcnh8 or Kcnh3 selectively alters nighttime firing rates in SCN neurons

To explore the roles of Kv12.1-/Kv12.2-encoded K^+^ channels in regulating spontaneous repetitive firing in the SCN, we obtained whole-cell current-clamp recordings, during the day (ZT7-ZT12) and at night (ZT19-ZT24), from SCN neurons in slices prepared from adult wild type (WT) mice and from animals harboring targeted disruptions in the *Kcnh8* (Kv12.1^−/−^) or *Kcnh3* (Kv12.2^−/−^) locus (Figure 1). Consistent with previous reports,^[7–10]^ these experiments revealed a large (*P* = 0.001, one-way ANOVA) day-night difference in the mean ± SEM repetitive firing rates of WT SCN neurons (Figure 1C and Table 1). In marked contrast, there were no measurable differences in the mean ± SEM daytime and nighttime repetitive firing rates of Kv12.1^−/−^ (*P* = 0.99, one-way ANOVA) or Kv12.2^−/−^ (*P* = 0.79, one-way ANOVA) SCN neurons (Figure 1C and Table 1). In addition, the loss of the day-night difference in the mean repetitive firing rates of Kv12.1^−/−^ and Kv12.2^−/−^ SCN neurons reflects a selective increase (compared with WT cells) in mean nighttime repetitive firing rates (Figure 1C and Table 1).

**Figure 1.**
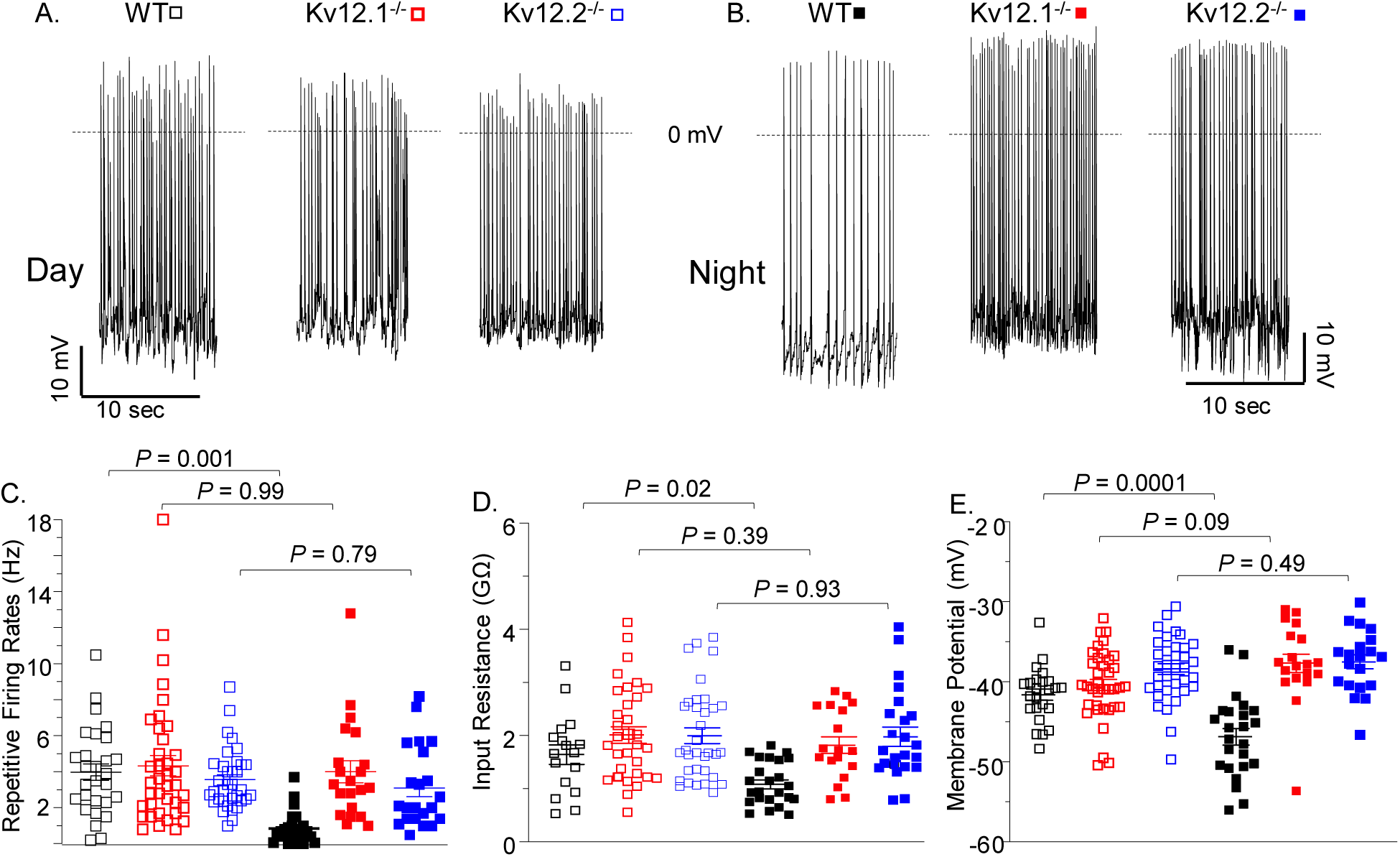
The repetitive firing rates of Kv12.1^−/−^ and Kv12.2^−/−^ SCN neurons are similar during the day and at night. Representative daytime (A) and nighttime (B) whole-cell current-clamp recordings from WT (□,▪), Kv12.1^−/−^ (□,▪) and Kv12.2^−/−^ (□,▪) SCN neurons are shown. (C) Daytime and nighttime repetitive firing rates, measured in individual WT, Kv12.1^−/−^, and Kv12.2^−/−^ SCN neurons, are plotted; mean ± SEM repetitive firing rates are indicated. As reported previously^7-10^, the mean ± SEM repetitive firing rate In WT SCN neurons is higher (*P* = 0.001, one-way ANOVA) during the day (n = 28) than at night (n = 25). In contrast, there are no day-night differences (one-way ANOVA) in the mean ± SEM repetitive firing rates of Kv12.1^−/−^ (*P* = 0.99; n = 21 - 38) or Kv12.2^−/−^ (*P* = 0.79; n = 22 - 35) SCN neurons. Input resistances (D) and membrane potentials (E), measured in individual WT, Kv12.1^−/−^, and Kv12.2^−/−^ SCN neurons during the day and at night, are plotted; mean ± SEM values and *P* values are indicated. In contrast to WT SCN neurons, there are no day-night differences (one-way ANOVA) in the mean ± SEM input resistances (D) or membrane potentials (E) of Kv12.1^−/−^ or Kv12.2^−/−^ SCN neurons.

**Table 1.** Resting and active membrane properties of WT, Kv12.1^−/−^, Kv12.2^−/−^ and DKO (Kv12.1^−/−^/Kv12.2^−/−^) SCN neurons during the day and at night*

|  |  | Firing rate<br>(Hz) | R<sub>in</sub><br>(GΩ) | V<sub>r</sub><br>(mV) | ISI<br>(ms) |
| --- | --- | --- | --- | --- | --- |
| Day ZT 7-12 | WT | 4.0 ± 0.5<sup>1</sup><br>n = 28 | 1.7 ± 0.2<sup>2</sup><br>n = 20 | -41.6 ± 0.7<sup>1</sup><br>n = 28 | 280 ± 9<sup>1</sup><br>n = 28 |
|  | Kv12.1<sup>-/-</sup> | 4.3 ± 0.6<br>n = 38 | 2.0 ± 0.2<br>n = 33 | -40.4 ± 0.7<br>n = 38 | 252 ± 9<br>n = 38 |
|  | Kv12.2<sup>-/-</sup> | 3.6 ± 0.3<br>n = 35 | 2.0 ± 0.1<br>n = 34 | -38.4 ± 0.7<br>n = 35 | 311 ± 9<br>n = 35 |
|  | DKO | 4.1 ± 0.3<br>n = 6 | 1.6 ± 0.2<br>n = 6 | -40.8 ± 1.6<br>n = 6 | 279 ± 13<br>n = 6 |
| Night ZT 18-24 | WT | 0.8 ± 0.2<sup>1</sup><br>n = 25 | 1.0 ± 0.1<sup>2</sup><br>n = 24 | -46.7 ± 1.0<sup>1</sup><br>n = 25 | 418 ± 29<sup>1</sup><br>n = 20 |
|  | Kv12.1<sup>-/-</sup> | 4.0 ± 0.7<br>n = 21 | 1.8 ± 0.2<br>n = 18 | -37.7 ± 1.2<br>n = 21 | 254 ± 6<br>n = 21 |
|  | Kv12.2<sup>-/-</sup> | 3.1 ± 0.5<br>n = 22 | 2.0 ± 0.2<br>n = 22 | -37.5 ± 0.9<br>n = 22 | 319 ± 17<br>n = 22 |
|  | DKO | 3.4 ± 0.4<br>n = 6 | 1.5 ± 0.2<br>n = 6 | -40.5 ± 3.2<br>n = 6 | 318 ± 12<br>n = 6 |

\*Values are means ± SEM; n = number of cells; R<sub>in</sub> = input resistance; V<sub>r</sub> = resting membrane potential; ISI = interspike interval. <sup>1,2</sup> Values in daytime and nighttime WT neurons are significantly (<sup>1</sup>P < 0.001; <sup>2</sup>P < 0.02) different.

Because the majority of the data (presented in Figure 1 and Table 1) was obtained during the latter portion of the night (i.e., ZT20-ZT24), additional recordings were obtained throughout the light-dark cycle. Specifically, additional cell-attached current-clamp recordings were obtained from cells in slices during the transition from ‘lights-on to lights-off’ (ZT12-ZT14), as well as slices prepared mid-day (ZT6-ZT8) and mid-evening (ZT18-ZT20). Analysis of these recordings revealed that during the transition from day to night, the mean repetitive firing rate of WT SCN neurons decreased to ∼ 1Hz (compared to ∼ 4 Hz at mid-day) and that this rate was maintained to the end of the night phase (Supplemental Figure 4). In marked contrast, the mean repetitive firing rates of Kv12.1^−/−^ and Kv12.2^−/−^ SCN neurons during the ‘day to night’ transition (∼3.5 Hz) were much higher (Supplemental Figure 4) and remained high at mid-day and mid-evening (Supplemental Figure 4). In addition, the mean ± SEM daytime repetitive firing rates of Kv12.1^−/−^ and Kv12.2^−/−^ SCN neurons are indistinguishable from WT SCN cells (Figure 1 C and Table 1). Similar results were obtained in recordings from DKO (Kv12.1^−/−^/Kv12.2^−/−^) SCN neurons lacking both Kv12.1 and Kv12.2 (Table 1). Consistent with the similarities in repetitive firing rates measured during the day and at night, Kv12.1^−/−^, Kv12.2^−/−^ and DKO SCN neurons do not display the day-night difference in interspike intervals characteristic of WT SCN neurons (Table 1). Taken together, these results demonstrate that the repetitive firing rates of Kv12.1^−/−^ and Kv12.2^−/−^ SCN neurons are increased throughout the night.

In contrast to WT cells, there were also no day-night differences in the mean ± SEM input resistances or resting membrane potentials of Kv12.1^−/−^, Kv12.2^−/−^ or DKO SCN neurons (Figure 1D, 1E and Table 1). Consistent with the repetitive firing data, these results reflect the higher mean input resistances and depolarized membrane potentials of nighttime Kv12.1^−/−^, Kv12.2^−/−^ and DKO SCN neurons, compared with nighttime WT SCN neurons (Figure 1D, 1E and Table 1). In contrast, the mean ± SEM daytime input resistances and resting membrane potentials of Kv12.1^−/−^, Kv12.2^−/−^ and DKO SCN neurons are similar to those measured in daytime WT SCN neurons (Figure 1D, 1E and Table 1).

### Acute in vivo knockdown of Kv12.1 or Kv12.2 also selectively increases nighttime firing rates

The electrophysiological studies above demonstrate increased repetitive firing rates and input resistances, reduced interspike intervals and depolarized resting membrane potentials in nighttime Kv12.1^−/−^, Kv12.2^−/−^ and DKO, compared with WT, SCN neurons; the repetitive firing and intrinsic membrane properties of daytime Kv12.1^−/−^, Kv12.2^−/−^, DKO and WT SCN neurons, in contrast, are indistinguishable (Figure 1 and Table 1). Interpreting the physiological significance and the functional implications of these findings is confounded, however, by the fact that the Kv12.1^−/−^, Kv12.2^−/−^ and DKO mice lack Kv12.1 or/and Kv12.2 throughout development. Additional experiments were undertaken, therefore, to test the hypotheses that Kv12.1 and/or Kv12.2 are required for the regulation of the nighttime repetitive firing rates and the membrane properties of *mature* SCN neurons using an interfering RNA^[31]^ strategy to knockdown the expression of the Kv12.1 and Kv12.2 subunits acutely in the SCN of adult animals *in vivo*.

To identify short hairpin RNA (shRNA) sequences that effectively reduce Kv12.1 (or Kv12.2) expression, five Kv12.1-targeted (or five Kv12.2-targeted) shRNAs were screened individually in tsA-201 cells co-transfected with a cDNA construct encoding Kv12.1-eYFP (or Kv12.2-eYFP). The Kv12.1-targeted (and Kv12.2-targeted shRNA) sequences providing the maximal (>90%) reductions in Kv12.1 (or Kv12.2) expression identified in these initial screens were further evaluated for specificity. In these experiments, tsA-201 cells were co-transfected with the validated Kv12.1-targeted or Kv12.2-targeted shRNA or a NT shRNA with cDNA constructs encoding Kv12.1-eYFP, Kv12.2-eYFP, or Kv4.1-eYFP (from another Kv subfamily). As illustrated in Figure 2, Western blot analysis revealed that the Kv12.1-targeted shRNA markedly reduced expression of Kv12.1-eYFP (Figure 2A), without measurably affecting Kv12.2-eYFP or Kv4.1-eYFP (Figure 2B and 2C). Similarly, the Kv12.2-targeted shRNA markedly reduced Kv12.2-eYFP, without measurably affecting Kv12.1-eYFP or Kv4.1-eYFP (Figure 2A-2C). These selective Kv12.1-targeted and Kv12.2-targeted shRNAs, as well as the NT shRNA, were then cloned (separately), in a microRNA (human miR30) context,^[32]^ into a plasmid containing a synapsin promoter and eGFP,^[19]^ and adeno-associated viruses serotype 8 (AAV8) were generated.

**Figure 2.**
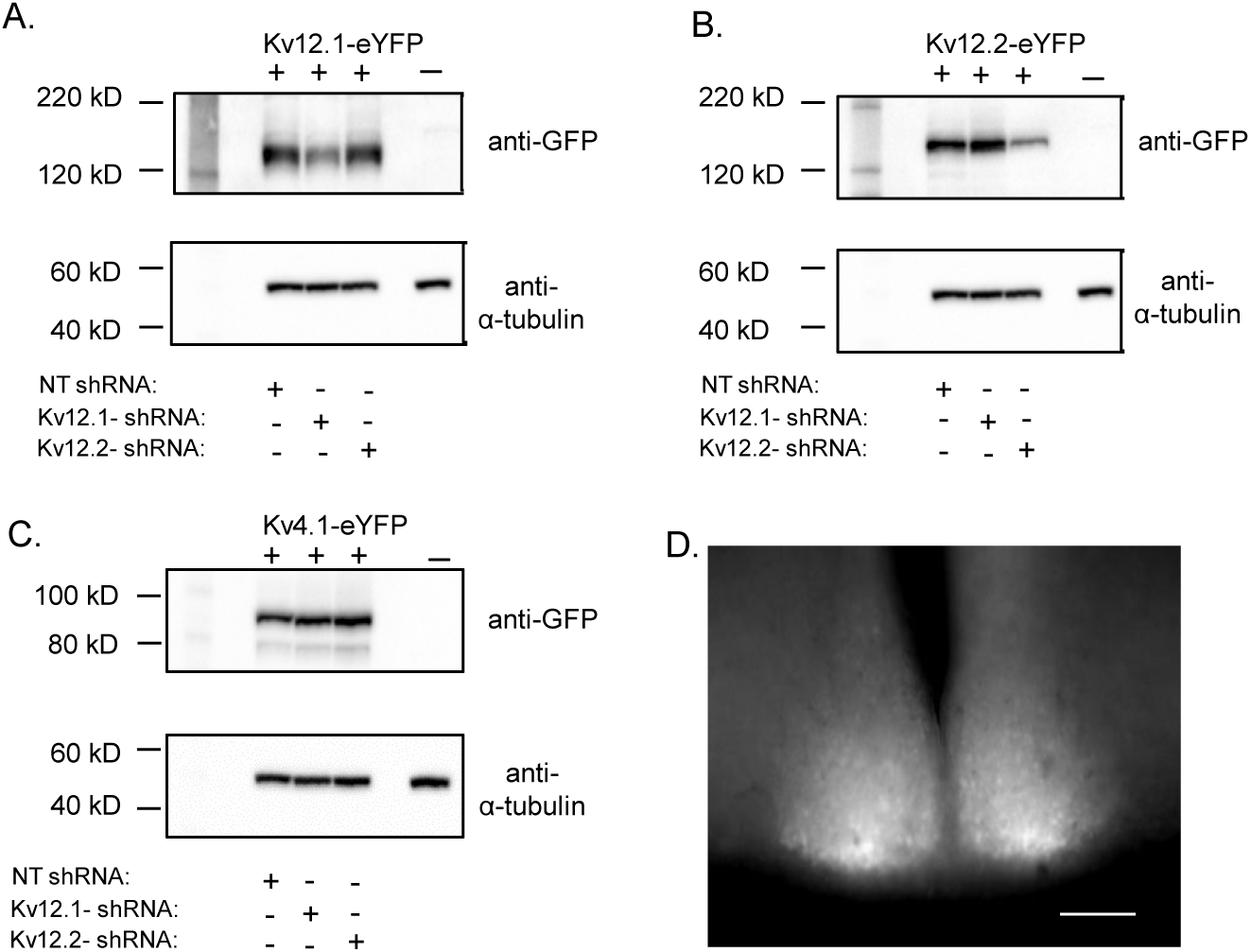
Validation of the Kv12.1-targeted and Kv12.2-targeted shRNAs. To examine the specificity of the selected Kv12.1-targeted and Kv12.2-targeted shRNAs, each was expressed in tsA-201 cells with a cDNA construct encoding Kv12.1-eYFP, Kv12.2-eYFP or Kv4.1-eYFP; parallel experiments were completed with the non-targeted (NT) shRNA. Approximately 24 hrs following transfections, cell lysates were prepared, fractionated by SDS-PAGE, transferred to PVDF membranes and probed with an anti-GFP antibody. All blots were also probed with an anti-α-tubulin antibody to verify equal protein loading. (A) Co-expression with the Kv12.1-targeted shRNA markedly reduced Kv12.1-eYFP protein expression, whereas the Kv12.2-targeted shRNA and the NT shRNA were without effects on Kv12.1-eYFP. (B) Similarly, co-expression with the Kv12.2-targeted shRNA, but not the Kv12.1-targeted or the NT shRNA, reduced Kv12.2-eYFP protein expression. (C) In contrast, neither the Kv12.1-targeted nor the Kv12.2-targeted shRNA measurably affected expression of Kv4.1-eYFP. (D) eGFP-expressing neurons were readily identified in acute SCN slices prepared two weeks following bilateral injections of the Kv12.1-targeted (or the Kv12.2-targeted) shRNA-expressing AAV8 into the SCN; scale bar = 250 μm.

Two weeks following stereotaxic injections of the Kv12.1-targeted, Kv12.2-targeted or NT shRNA-expressing AAV8 into the adult SCN, whole-cell current-clamp recordings were obtained from eGFP-positive SCN neurons in slices (Figure 2D). Recordings from NT shRNA-expressing SCN neurons during the daytime (ZT7-ZT12) and nighttime (ZT18-ZT24) revealed a day-night difference in mean ± SEM repetitive firing rates (Figure 3C and Table 2). Similar to WT neurons (Table 1), the mean ± SEM repetitive firing rate measured in NT shRNA-expressing SCN neurons was much (*P = 0.0001*, one-way ANOVA) higher during the day than at night (Figure 3C and Table 2). The mean ± SEM repetitive firing rates of daytime Kv12.1-targeted shRNA- (*P* = 0.39) and Kv12.2-targeted (*P* = 0.74) shRNA-expressing SCN neurons were similar to those measured in daytime NT shRNA-expressing neurons (One-way ANOVA; Figure 3C and Table 2). In contrast, the mean ± SEM repetitive firing rates of nighttime Kv12.1-targeted (*P* = 0.0001) and Kv12.2-targeted (*P* = 0.007) shRNA-expressing SCN neurons were much higher than in nighttime NT shRNA-expressing SCN neurons (One-way ANOVA; Figure 3C and Table 2). In addition, as is also evident in Figure 3C, the day-night difference in repetitive firing rate, evident in NT shRNA-expressing SCN neurons, was not observed in Kv12.1-targeted or Kv12.2-targeted shRNA-expressing (Table 2) SCN neurons.

**Figure 3.**
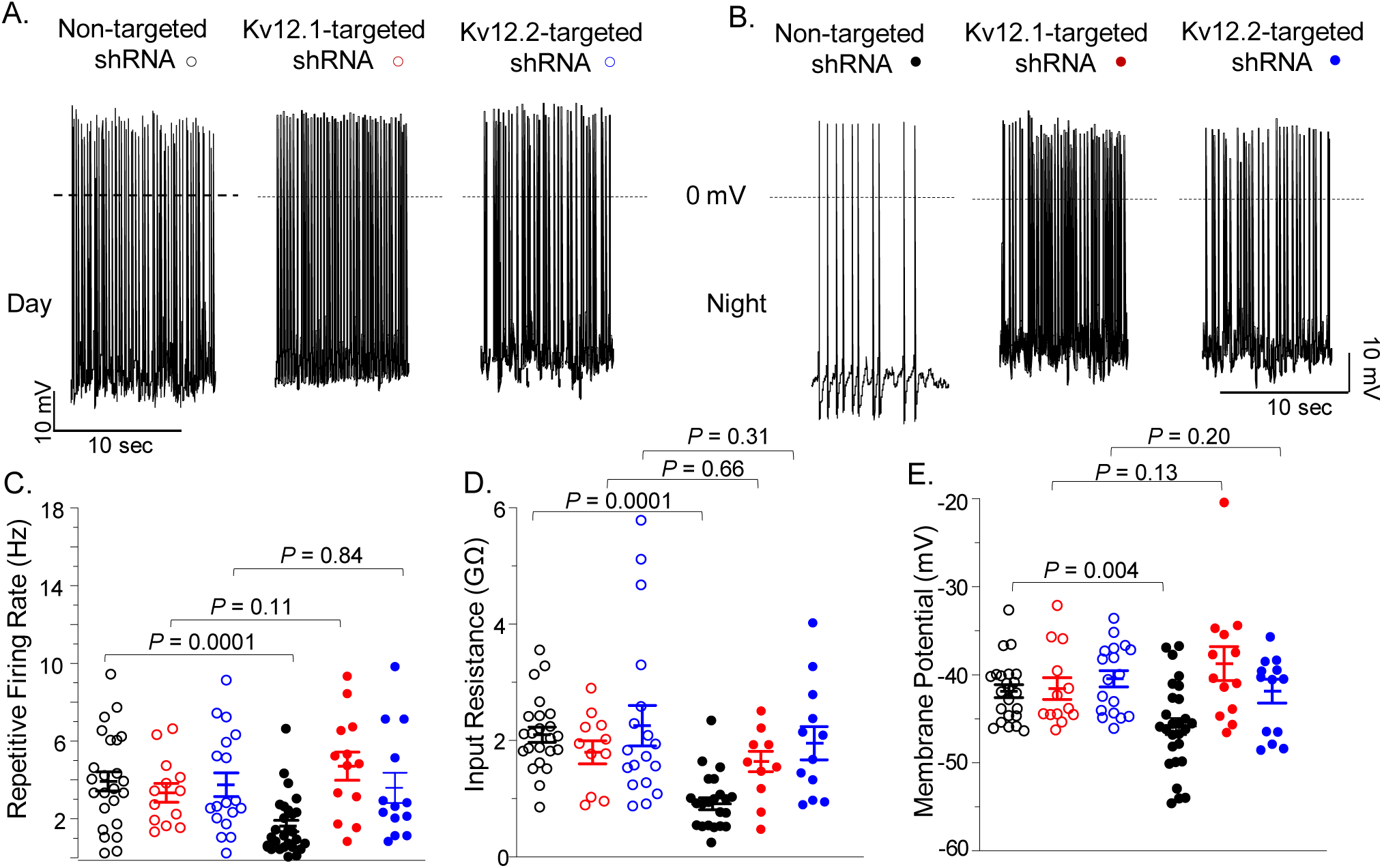
Acute *in vivo* shRNA-mediated knockdown of Kv12.1 or Kv12.2 increases the nighttime, but *not* the daytime, repetitive firing rates of SCN neurons. Representative daytime (A) and nighttime (B) whole-cell current-clamp recordings, obtained from NT shRNA- (○,●), Kv12.1-targeted shRNA- (○,●), and Kv12.2-targeted shRNA- (○,●) expressing SCN neurons during the day (ZT7-ZT12) and at night (ZT18-ZT24), are shown. (C) Daytime and nighttime repetitive firing rates, measured in individual SCN neurons expressing the NT, Kv12.1-targeted, or Kv12.2-targeted shRNA, are plotted; mean ± SEM repetitive firing rates are indicated. Similar to WT SCN^[7–10]^ neurons (Figure 1), the mean ± SEM repetitive firing rate of NT shRNA-expressing SCN neurons was lower (*P* = 0.0001, one-way ANOVA) at night (n = 25) than during the day (n = 24). In marked contrast, the mean ± SEM repetitive firing rates of Kv12.1- and Kv12.2-targeted shRNA-expressing SCN neurons at night are not (one-way ANOVA) different from daytime firing rates. The input resistances (D) and membrane potentials (E) of NT shRNA-, Kv12.1-targeted shRNA-, and Kv12.2-targeted shRNA-expressing SCN neurons during the day and at night were also determined. Similar to WT SCN neurons,^[7–10]^ the mean ± SEM input resistance of NT shRNA-expressing SCN neurons was higher (*P* = 0.0001, one-way ANOVA), during the day than at night, and the mean ± SEM membrane potential was more depolarized (*P* = 0.004, one-way ANOVA), during the day than at night. In contrast, there were no day-night differences (one-way ANOVA) in the mean ± SEM input resistances (D) or membrane potentials (E) of Kv12.1-targeted shRNA- and Kv12.2-targeted shRNA-expressing SCN neurons.

**Table 2.** Resting and active membrane properties of non-targeted- (NT), Kv12.1-targeted and Kv12.2-targeted shRNA-expressing SCN neurons during the day and at night*

| | | Firing rate<br>(Hz) | $R_{in}$<br>(GΩ) | $V_r$<br>(mV) | ISI<br>(ms) |
| --- | --- | --- | --- | --- | --- |
| Day ZT 7-12 | Non-targeted<br>shRNA | $3.9 \pm 0.5$<sup>1</sup><br>n = 24 | $2.1 \pm 0.1$<sup>1</sup><br>n = 23 | $-41.5 \pm 0.7$<sup>2</sup><br>n = 24 | $221 \pm 4$<sup>1</sup><br>n = 24 |
| | Kv12.1-targeted<br>shRNA | $3.3 \pm 0.5$<br>n = 13 | $1.8 \pm 0.2$<br>n = 11 | $-41.5 \pm 1.2$<br>n = 13 | $213 \pm 9$<br>n = 13 |
| | Kv12.2-targeted<br>shRNA | $3.8 \pm 0.6$<br>n = 18 | $2.3 \pm 0.3$<br>n = 18 | $-40.5 \pm 0.9$<br>n = 18 | $243 \pm 8$<br>n = 18 |
| Night ZT 18-24 | Non-targeted<br>shRNA | $1.6 \pm 0.3$<sup>1</sup><br>n = 27 | $0.9 \pm 0.1$<sup>1</sup><br>n = 22 | $-45.8 \pm 1.0$<sup>2</sup><br>n = 27 | $366 \pm 24$<sup>1</sup><br>n = 27 |
| | Kv12.1-targeted<br>shRNA | $4.7 \pm 0.7$<br>n = 13 | $1.6 \pm 0.2$<br>n = 9 | $-38.7 \pm 1.9$<br>n = 13 | $187 \pm 5$<br>n = 13 |
| | Kv12.2-targeted<br>shRNA | $3.6 \pm 0.8$<br>n = 13 | $1.9 \pm 0.3$<br>n = 12 | $-42.6 \pm 1.5$<br>n = 13 | $234 \pm 6$<br>n = 13 |

\*Values are means ± SEM; n = number of cells; $R_{in}$ = input resistance; $V_r$ = resting membrane potential; ISI = Interspike interval. <sup>1,2</sup> Values in daytime and nighttime NT shRNA-expressing neurons are significantly (<sup>1</sup> $P < 0.001$ ; <sup>2</sup> $P < 0.005$ ) different.

Similar to WT SCN neurons (Table 1), input resistances were higher (*P* = 0.02, one-way ANOVA), resting membrane potentials were more depolarized (*P* = 0.0001, one-way ANOVA) and interspike intervals were shorter (*P* = 0.0001, one-way ANOVA) in NT shRNA-expressing SCN neurons during the day than at night (Figure 3D, 3E and Table 2). In contrast, the input resistances, resting membrane potentials and interspike intervals measured in Kv12.1-targeted and Kv12.2-targeted shRNA-expressing SCN neurons (*P* = 0.93, one-way ANOVA) at night and during the day are similar (Figure 3D, Figure 3E and Table 2). The functional effects of the acute *in vivo* knockdown of Kv12.1 or Kv12.2 (Table 2) on the repetitive firing rates and membrane properties of SCN neurons, therefore, are virtually identical to those observed in Kv12.1^−/−^ and Kv12.2^−/−^ (Table 1) SCN neurons (see: **Discussion**).

### Kv12-encoded current densities are higher in nighttime, than in daytime, SCN neurons

The selective effects of the targeted deletion or the acute knockdown of Kv12.1 or Kv12.2 on the nighttime repetitive firing rates and membrane properties of SCN neurons suggest day-night differences in the functional expression of Kv12-encoded K^+^ currents. To explore this hypothesis directly, whole-cell voltage-clamp recordings were obtained from daytime and nighttime WT and DKO (Kv12.1^−/−^/Kv12.2^−/−^) SCN neurons before and after application of 20 μM CX4, a selective blocker of heterologously expressed Kv12.2 channels.^[30]^ In these experiments, whole-cell outward K^+^ currents, evoked in response to action potentials waveforms (Figure 4A) obtained from nighttime WT SCN neurons, were recorded from daytime and nighttime WT (Figure 4B) and DKO (Figure 4C) SCN neurons with TTX and CdCl_2_ in the bath to block voltage-gated Na^+^ and Ca^2+^ channels, respectively. Offline digital subtraction of the currents recorded before and after exposure to CX4 provided the CX4-sensitive outward K^+^ currents. As is evident in the representative recordings shown in Figure 4B, the density of the CX4-sensitive K^+^ current is higher in the nighttime, compared with the daytime, WT SCN neuron. Similar results were obtained in recordings from multiple WT neurons (Figure 4D). The mean ± SEM CX4-sensitive current densities measured in daytime and nighttime WT SCN neurons were 7.9 ± 1.8 pA/pF (n = 17) and 23.6 ± 6.3 pA/pF (n = 13), respectively. The marked day-night difference in the densities of the CX4-sensitive outward K^+^ currents observed in WT SCN neurons, however, is not evident in DKO SCN neurons (Figure 4C). The mean ± SEM CX4-sensitive outward K^+^ current densities measured in daytime and nighttime DKO SCN neurons were statistically indistinguishable at 4.3 ± 1.2 pA/pF (n = 8) and 5.6 ±1.3 pA/pF (n = 8), respectively (Figure 4D). The densities of the CX4-sensitive outward K^+^ currents in daytime and nighttime DKO SCN neurons, therefore, are quite similar.

**Figure 4.**
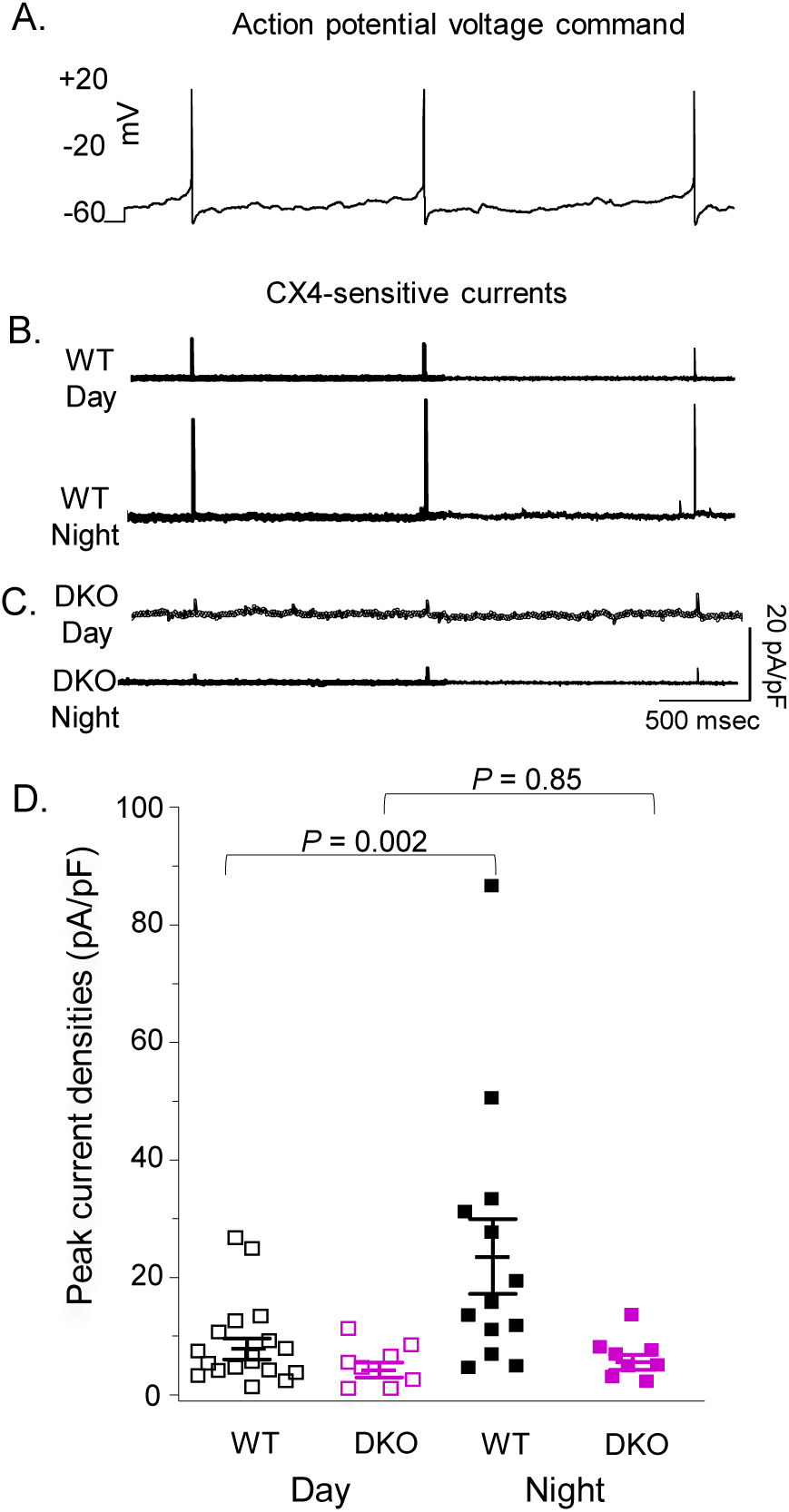
CX4-sensitive K^+^ current densities in SCN neurons are higher at night than during the day. (A) Whole-cell outward K^+^ currents, evoked by an action potential voltage command recorded from a nighttime WT SCN neuron, were recorded in WT and DKO (Kv12.1^−/−^/Kv12.2^−/−^) SCN neurons before and after local application 20 μM CX4. Subtraction of the currents recorded before/after application of CX4 in each cell provided the CX4-sensitive K^+^ currents. Representative CX4-sensitive outward K^+^ currents, obtained by offline digital subtraction of the currents recorded in the presence of CX4 from the currents recorded (in the same cell) in the control bath solution during the day and at night in WT (B) and DKO (C) SCN neurons, are shown. (D) Peak CX4-sensitive K^+^ current densities, measured in WT (□,▪) and DKO (□,▪) SCN neurons during the day and at night, are plotted. The mean ± SEM CX4-sensitive K^+^ current density in WT SCN neurons is much higher (*P =* 0.002, one-way ANOVA) at night (n = 13) than during the day (n = 17). In contrast, CX4-sensitive K^+^ current densities in DKO SCN neurons are low during the day (n = 8) and at night (n = 8), and there is no day-night difference (*P* = 0.85, one-way ANOVA) in mean ± SEM CX4-sensitive K^+^ current densities in DKO SCN neurons.

Although the action potential- (voltage-) clamp recordings (Figure 4) reveal that the mean ± SEM CX4-sensitive outward K^+^ current density is considerably higher in nighttime, than in daytime, WT SCN neurons, there were a couple of daytime WT SCN neurons with higher than average CX4-sensitive current densities. In addition, the CX4-sensitive current densities measured in individual nighttime WT SCN neurons were quite variable (Figure 4D). To explore the functional consequences of these apparent differences, the effects of CX4 on the repetitive firing properties of SCN neurons were examined in whole-cell current-clamp recordings from cells in acute slices prepared from WT animals during the day (ZT7-ZT12) or at night (ZT18-ZT24). Consistent with the action potential- (voltage-) clamp data (Figure 4), these experiments revealed that application of 20 μM CX4 increased repetitive firing rates in only a small fraction (2 of 17; ∼10%) of daytime WT SCN neurons (Figure 5C), whereas the repetitive firing rates of most (11 of 15; ∼70%) nighttime WT SCN neurons were increased following application of CX4 (Figure 5D). In addition to confirming day-night differences in the functional expression of Kv12-encoded K^+^ currents, these observations suggest that there is cellular heterogeneity in the subthreshold K^+^ channels mediating the day-night switch in the repetitive firing rates of SCN neurons (see **Discussion**).

**Figure 5.**
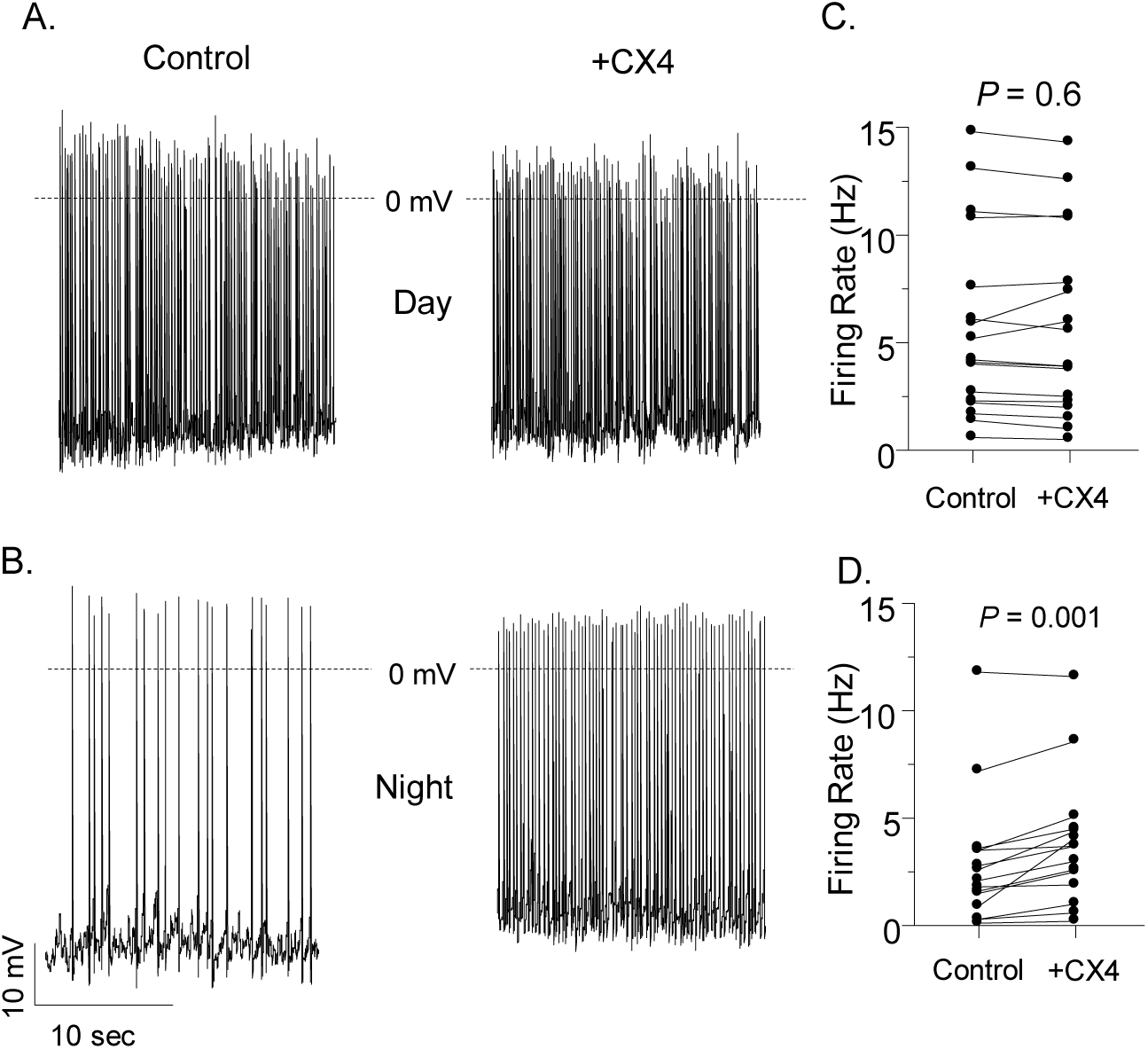
Pairwise comparisons of repetitive firing rates in WT SCN neurons in the absence and presence of CX4. Representative daytime (A) and nighttime (B) whole-cell current-clamp recordings obtained from WT SCN neurons before and after application of 20 μM CX4 are shown. The repetitive firing rates measured in individual daytime (C) and nighttime (D) WT SCN neurons before and after application of 20 μM CX4 are plotted. In the vast majority (15 of 17; ∼90%) of WT SCN neurons, daytime repetitive firing rates were not affected (*P =* 0.6, repeated measures t-test) by the application of 20 μM CX4 (C). In contrast, nighttime repetitive firing rates were increased (*P =* 0.001, repeated-measures Student t-test) in most (11 of 15; ∼70%) WT SCN neurons upon application of 20 μM CX4 (D). Approximately 30% (4 of 15) of nighttime WT SCN neurons, however, were not measurably affected by 20 μM CX4 (see text).

### Dynamic clamp-mediated subtraction of I_Kv12_ increases firing rate in nighttime WT SCN neurons

To explore the impact of changing Kv12-encoded current amplitudes on the spontaneous repetitive firing rates of SCN neurons directly, we employed dynamic clamp to manipulate the Kv12-encoded current (I_Kv12_) electronically in real time during current-clamp recordings. As described in **Materials and Methods**, a model of I_Kv12_ was developed using published data on heterologously expressed Kv12.1 currents.^[24]^ The steady-state voltage-dependence of activation of I_Kv12_ in the model was then adjusted to fit the acquired voltage- (Supplemental Figure 2A, 2B) and action potential- (Figure 4) clamp data from WT SCN neurons. The properties and amplitudes of I_Kv12_ produced by the model (Supplemental Figure 2C) reliably reproduce the CX4-sensitive currents identified in action potential-clamp recordings from WT SCN neurons.

In initial dynamic clamp experiments, whole-cell current-clamp recordings were obtained from SCN neurons in acute slices prepared from adult WT animals at night or during the day. After recording gap-free spontaneous firing activity under control conditions (Figure 6A, 6E), the effects on spontaneous repetitive firing rates of subtracting increasing I_Kv12_ amplitudes, presented in multiples of the minimal I_Kv12_ amplitude, x (determined as described in Materials and Methods, were determined. As illustrated in the representative recordings from a WT nighttime SCN neuron in Figure 6 A-C, subtraction of the minimal I_Kv12_ (-x) increased the rate of repetitive firing (Figure 6B), and the rate was increased further when the amplitude of I_Kv12_ was increased (-3x) threefold (Figure 6C). In contrast, as illustrated in the representative recordings from a WT daytime SCN neuron in Figure 6E-G, dynamic clamp-mediated subtraction of I_Kv12_ (at -x or -3x) had little to no effect on the rate of spontaneous action potential firing. Similar results were obtained in recordings from additional WT nighttime and daytime SCN neurons in which varying amplitudes (-x, -2x, -3x, -4x, and -5x) of I_Kv12_ were subtracted via dynamic clamp. The mean ± SEM percent changes in the spontaneous repetitive firing rates of WT daytime (□; n = 7) and nighttime (▪; n = 10) SCN neurons are plotted as a function of the magnitude of I_Kv12_ subtracted in Figure 6, panels D and H. The *Insets* below panels C and G in Figure 6 show the waveforms of individual action potentials recorded in the representative WT nighttime (C) and daytime (G) SCN neurons, plotted on an expanded timescale. The modeled (-3x) I_Kv12_ waveforms generated in these cells are shown below the voltage records; the zero current levels are indicated by dotted lines. As evident, the mean peak I_Kv12_ amplitudes, revealed during action potentials, are substantially higher in the nighttime (Figure 6C, *Inset*), than in the daytime (Figure 6G, *Inset*), SCN neuron. In addition, in the nighttime neuron, (Figure 6C, Inset), I_Kv12_ increases as the membrane potential approaches the threshold for generating an action potential. Furthermore, the mean modeled I_Kv12_ amplitudes are markedly larger during interspike intervals in nighttime (1.49 ± 0.15 pA, Figure 6C, *Inset*), than in the daytime (-0.39 ± 0.05 pA, Figure 6G, *Inset*), SCN neurons. These are observations consistent with a selective role for I_Kv12_ in regulating the interspike interval and the rate of spontaneous repetitive action potential firing in nighttime SCN neurons.

**Figure 6.**
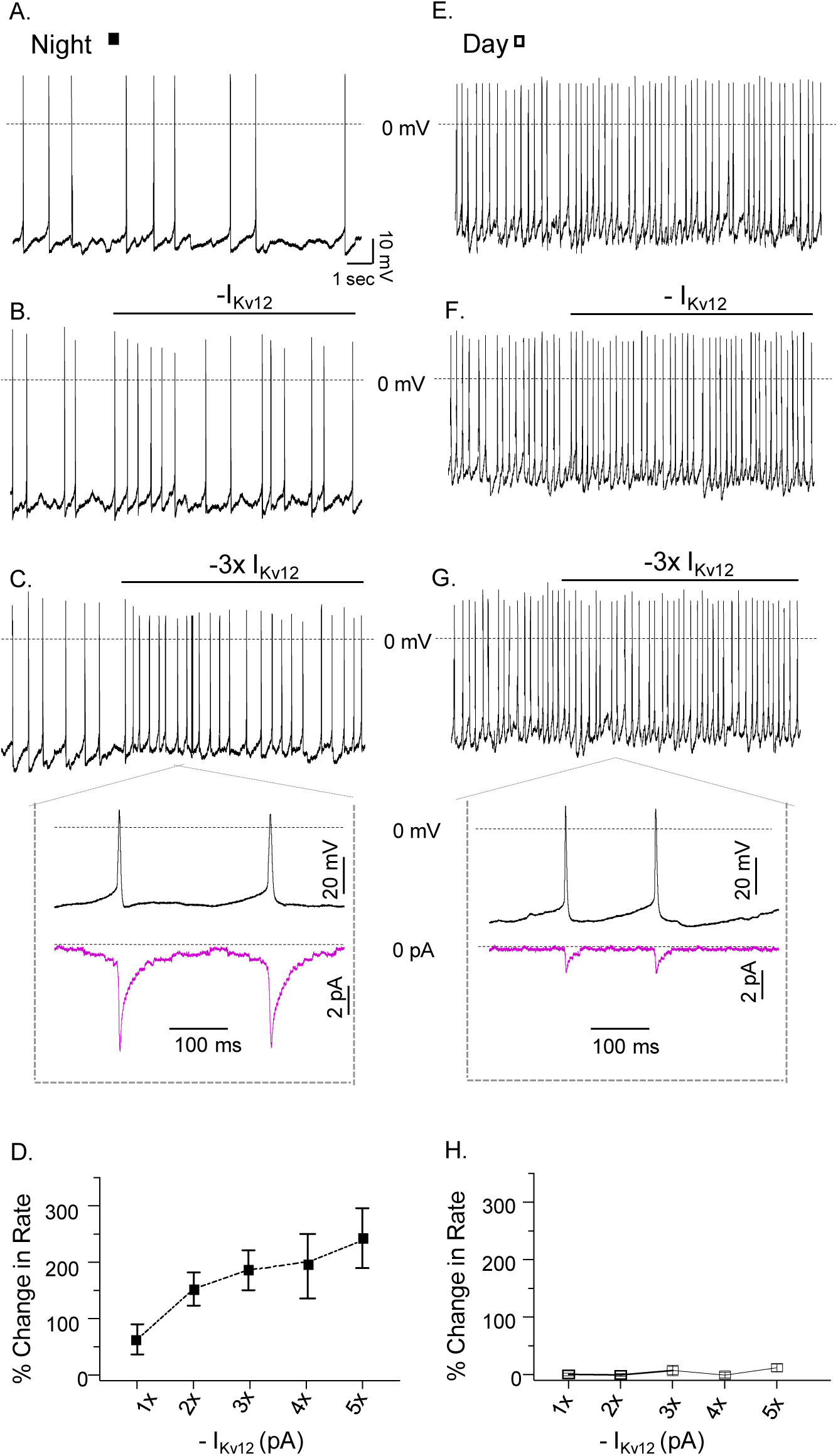
Dynamic clamp-mediated subtraction of I_Kv12_ increases the repetitive firing rates of nighttime WT SCN neurons. Representative whole-cell current-clamp recordings from a WT nighttime (A-C) and a WT daytime (E-G) SCN neuron at baseline (A,E) and with different magnitudes of modeled I_Kv12_ subtracted (B-C, F-G) via dynamic clamp are shown. In the *insets* below panels C and G, the waveforms of individual action potentials (black) recorded in the representative WT nighttime (C) and daytime (G) SCN neurons are displayed, plotted on an expanded timescale. Below these voltage records, the modeled (-3x) I_Kv12_ waveforms (purple) are shown; the zero current levels are indicated by the dotted lines. As is evident, the effect of subtracting increasing I_Kv12_ is greater in the nighttime (B,C), than in the daytime (F,G), WT SCN neuron. Similar results were obtained in current-clamp recordings from additional WT nighttime and daytime SCN neurons in which varying amplitudes (-x, -2x, -3x, -4x, and -5x) of the minimal modeled I_Kv12_ (2 pA) were subtracted via dynamic clamp. (D,H) The mean ± SEM percent changes in the spontaneous repetitive firing rates of WT daytime (□; n = 10) and nighttime (▪; n = 7) SCN neurons are plotted as a function of the magnitude of modeled I_Kv12_ subtracted.

In contrast with the marked differences in the effects of subtracting I_Kv12_ in WT nighttime and daytime SCN neurons (Figure 6), dynamic clamp-mediated addition of I_Kv12_ decreased the repetitive firing rates of both nighttime and daytime WT SCN neurons (Figure 7). Addition of the modeled I_Kv12_ also reduced the rate of repetitive firing in nighttime Kv12.1^−/−^ SCN neurons (Supplemental Figure 5A-C), although, with small current injections, the magnitudes of the percent changes in repetitive firing rates in nighttime Kv12.1^−/−^ SCN neurons were smaller than in nighttime WT SCN neurons (Supplemental Figure 5D).

**Figure 7.**
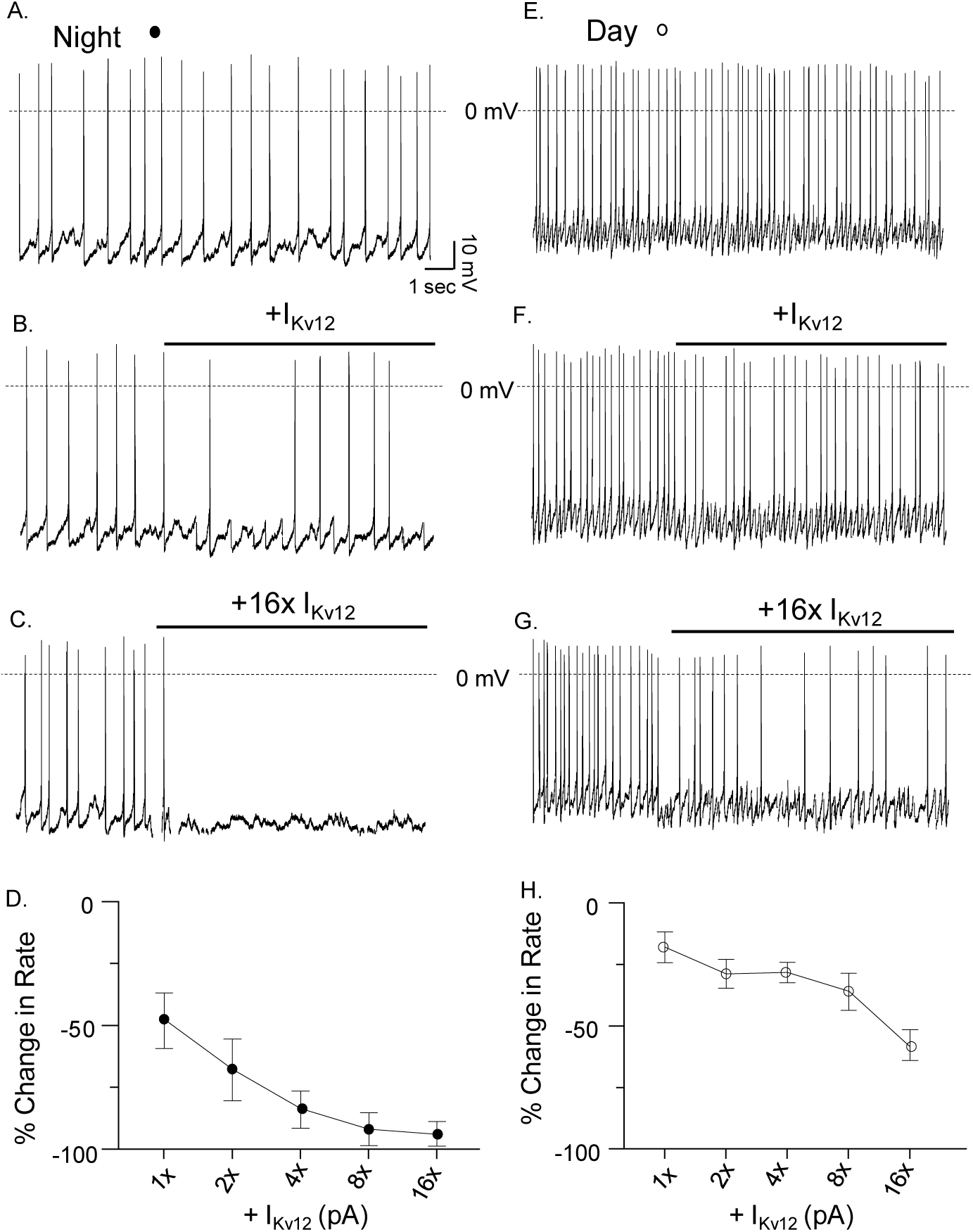
Dynamic clamp-mediated addition of modeled Kv12-encoded (I_Kv12_) currents in WT SCN neurons. Representative whole-cell current-clamp recordings from a WT nighttime (A-C) and a WT daytime (E-G) SCN neuron at baseline (A,E) and with different magnitudes of modeled I_Kv12_ added (B-C, F-G) via dynamic clamp are shown. As is evident, increasing I_Kv12_ reduced the rate of repetitive firing in both the WT nighttime (B,C) and the WT daytime (F,G) SCN neuron, although the impact is greater on nighttime (B,C), than on daytime (F,G), firing. Similar results were obtained in recordings from additional WT nighttime and daytime SCN neurons in which varying amplitudes (+x, +2x, +4x, +8x, and +16x) of the minimal modeled I_Kv12_ (0.5 pA) were added during current-clamp recordings. (D,H) The mean ± SEM percent changes in the spontaneous repetitive firing rates of WT daytime (○, n = 7) and nighttime (●, n = 10) SCN neurons are plotted as a function of the magnitude of modeled I_Kv12_ added.

To determine if the functional effects of dynamic clamp-mediated manipulation of I_Kv12_ observed in WT SCN neurons (Figures 6 and 7) are specific for I_Kv12_ or, alternatively, would be expected if any subthreshold voltage-activated K^+^ current were added/subtracted, we generated a mathematical model of I_A_, in SCN neurons. In previous studies, we have shown that shRNA-mediated knockdown of Kv4.1 decreases I_A_ amplitudes and increases the repetitive firing rates of daytime *and* nighttime SCN neurons.^[19]^ Using acquired voltage-clamp data describing the time- and voltage-dependent properties of I_A_ in WT SCN neurons,^[19]^ we generated a model of I_A_ gating (Supplemental Figure 3). In experiments similar to those described above for I_Kv12_, we determined the effects on spontaneous repetitive rates of subtracting/adding I_A_, presented in multiples of the minimal I_A_, x, determined as described in Materials and Methods. Dynamic clamp-mediated subtraction of modeled I_A_ initially increased the repetitive firing rates of WT nighttime (Figure 8A,C,E) and daytime (Figure 8B,D,F) SCN neurons. With the subtraction of higher amplitude currents, however, repetitive firing rates decreased in WT nighttime (Figure 8E) and daytime (Figure 8F) SCN neurons. The *Insets* below panels C and D in Figure 8 show the waveforms of individual action potentials recorded in the representative WT nighttime (C) and daytime (D) SCN neurons, plotted on an expanded timescale. The modeled (-3x) I_A_ waveforms generated in these cells are shown below the voltage records: the zero current levels are indicated by dotted lines. As is evident, there are clear differences in the peak amplitudes of the modeled I_A_ during the action potentials in WT nighttime (C) and daytime (D) SCN neurons. In addition and, in contrast with the results obtained with I_Kv12_ (Figure 6), I_A_ is prominent during interspike intervals in WT nighttime (-6.88 ± 0.83 pA, Figure 8C, *Inset*) and daytime (-7.03 ± 0.34 pA, Figure 8D, *Inset*) SCN neurons, an observation consistent with the physiological role of I_A_ driving repetitive firing rates in SCN neurons during the day and night.^[19]^ The addition of modeled I_A_ also decreased spontaneous repetitive firing rates similarly in WT nighttime (Figure 9A,C,E) and daytime (Figure 9B,D,F) SCN neurons.

**Figure 8.**
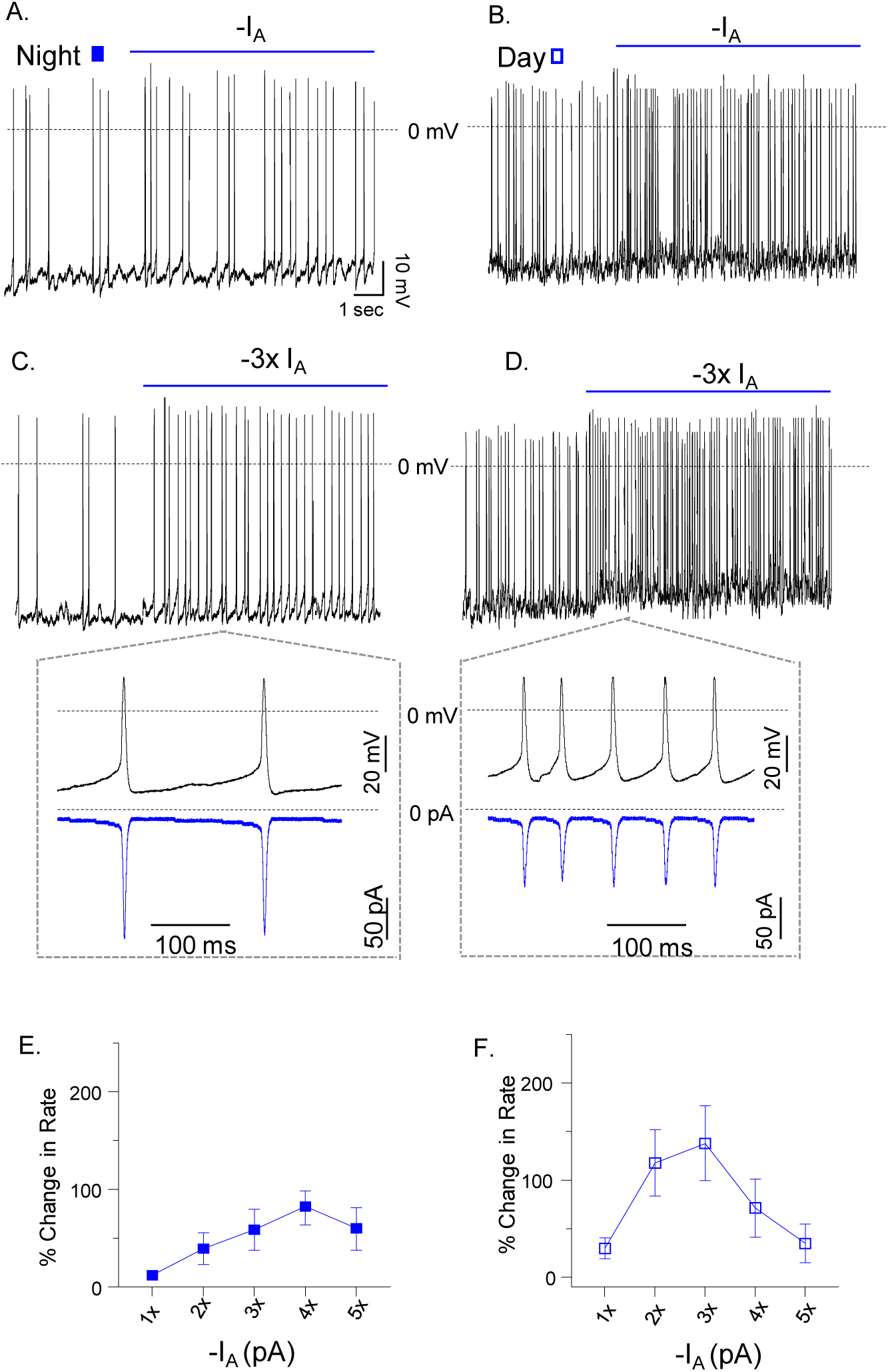
Dynamic clamp-mediated subtraction of modeled I_A_ in WT SCN neurons. Representative whole-cell current-clamp recordings from WT nighttime (A, C) and daytime (B, D) SCN neurons with different magnitudes of modeled I_A_ subtracted are shown. In the *insets* below panels C and D, the waveforms of individual action potentials (black), recorded in the representative WT nighttime (C) and daytime (D) SCN neurons, are plotted on an expanded timescale. The modeled (-3x) I_Kv12_ waveforms (blue) generated in these cells are shown; the zero current levels are indicated by the dotted lines. As is evident, the rates of repetitive firing of the representative nighttime (A,C) and daytime (B,D) SCN neurons are increased with the subtraction of modeled I_A_. Similar results were obtained in recordings from additional nighttime and daytime WT SCN in which varying amplitudes of the minimal modeled I_A_ (20 pA) were subtracted (-x, -2x, -3x, -4x, -5x) during current-clamp recordings. (E-F) The mean ± SEM percent changes in the spontaneous repetitive firing rates of WT SCN neurons in response to subtracting (□, □ = 16) modeled I_A_ are plotted as a function of the magnitude of the modeled I_A_ subtracted.

**Figure 9.**
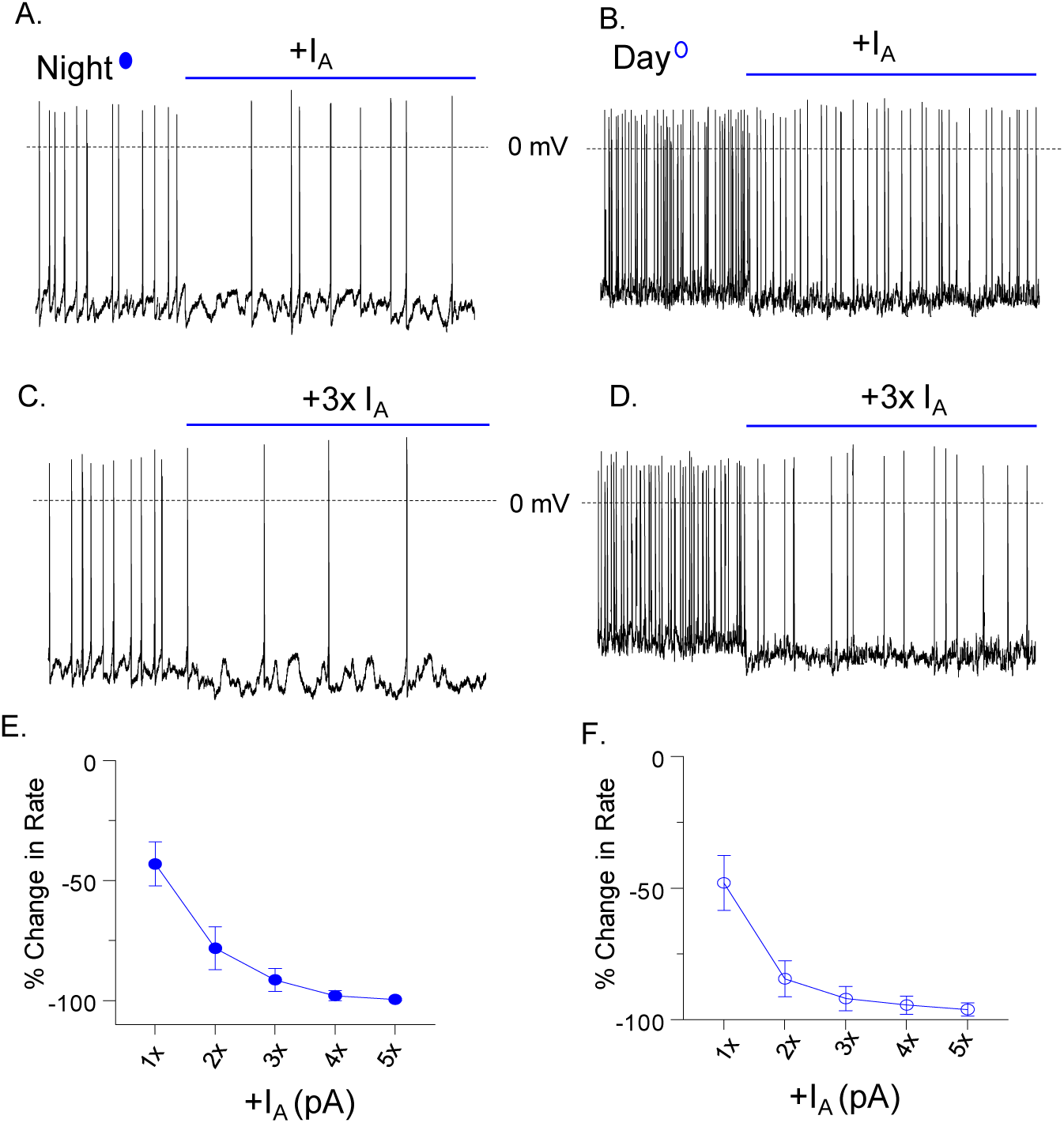
Dynamic clamp-mediated addition of modeled I_A_ in WT SCN neurons. Representative whole-cell current-clamp recordings from WT nighttime (A, C) and daytime (B, D) SCN neurons with different magnitudes of modeled I_A_ added are shown. The rates of repetitive firing of *both* the representative nighttime (A,C) and the representative daytime (B,D) SCN neuron are decreased with the addition of modeled I_A_. Recordings from additional nighttime and daytime WT SCN neurons in which varying amplitudes (x, 2x, 3x, 4x, 5x) of the minimal modeled I_A_ (20 pA) were added also revealed a decrease in repetitive firing rates. (E-H) The mean ± SEM percent changes in the spontaneous repetitive firing rates of WT SCN neurons in response to adding •, • = 14) modeled I_A_ are plotted as a function of the magnitude of the modeled I_A_ added.

Additional experiments were also completed to examined the effects of subtracting modeled I_Kv12_ versus modeled I_A_ in the same daytime cell. As in the experiments presented in Figure 8, the subtraction of I_A_ initially increased repetitive firing rates (Supplemental Figure 6A-B), and at higher current amplitudes, firing became unreliable (Supplemental Figure 6C). In contrast, subtraction of I_Kv12_ in the same cell had very little effect on repetitive firing rates (Supplemental Figure 6D-F), results similar to those presented in Figure 6. The functional effects of subtracting modeled I_A_ versus modeled I_Kv12_ on the repetitive firing rates of SCN neurons, therefore, are distinct.

### Kcnh8 and Kcnh3 transcript expression in the SCN do not display 24hr rhythms

The results of the action potential- (voltage-) clamp experiments (Figure 4) demonstrate that amplitudes/densities of the CX4-sensitive, Kv12-encoded K^+^ currents in SCN neurons are higher at night than during the day suggesting that the functional expression of the underlying K^+^ channels is regulated by the molecular clock. To determine if this regulation reflects day-night differences in the expression levels of the *Kcnh* transcripts encoding the Kv12.1 (*Kcnh8*) and Kv12.2 (*Kcnh3*) α subunits, quantitative real-time PCR (qRT-PCR) analysis was performed on RNA samples extracted from WT SCN (n = 7 - 8) every 4 hrs over 2 consecutive days. The expression levels of the transcripts encoding the clock genes, *Per2* and *Bmal*,^[1–4]^ and of *Hprt* (control gene) in every sample were also determined. Expression of the *Per2*, *Bmal*, *Kcnh8* and *Kcnh3* transcripts was normalized to *Hprt* in the same sample and evaluated for rhythmicity using JTK cycle analysis^[46]^ with the period set to 24 hr. As expected, these analyses revealed that *Per2* and *Bmal* display ∼ 24 hr anti-phase rhythms (Supplemental Figure 7) in expression.^[1–4]^ In contrast, the mRNA expression levels of *Kcnh8* and *Kcnh3* do not (*P* > 0.05, JTK cycle Bonferroni-adjusted) display 24 hr rhythms (Supplemental Figure 7), indicating that oscillations in the expression of the *Kcnh8* and/or *Kcnh3* transcripts do not underlie the observed day-night differences in the functional expression of Kv12-encoded currents (see: **Discussion**).

### Loss of Kv12 channels does not eliminate rhythms in (wheel-running) locomotor activity

Representative recordings of wheel-running activity in WT, Kv12.1^−/−^, and Kv12.2^−/−^ mice are presented in Figure 9A. In these experiments, all mice were entrained to a standard 12:12 hr LD cycle for 10 days, followed by at least 20 days in DD (Figure 10A). Circadian periods were measured in each animal after 20 days in DD (Figure 10C). The mean ± SEM circadian periods determined in adult WT (23.6 ± 0.1 hr; n = 9), Kv12.1^−/−^ (23.5 ± 0.1 hr; n = 15), Kv12.2^−/−^ (23.6 ± 0.1 hr; n = 14), and DKO (23.4 ± 0.1 hr; n = 12, *data not shown*) animals are indistinguishable (Figure 9C). Additional experiments, conducted on WT mice two weeks following bilateral injections of the Kv12.1-targeted, Kv12.2-targeted or NT-shRNA-expressing AAV8 into the SCN, revealed that the acute *in vivo* knockdown of Kv12.1 or Kv12.2 in the adult SCN also did not disrupt rhythmic wheel-running activity (Figure 10B and 10D).

**Figure 10.**
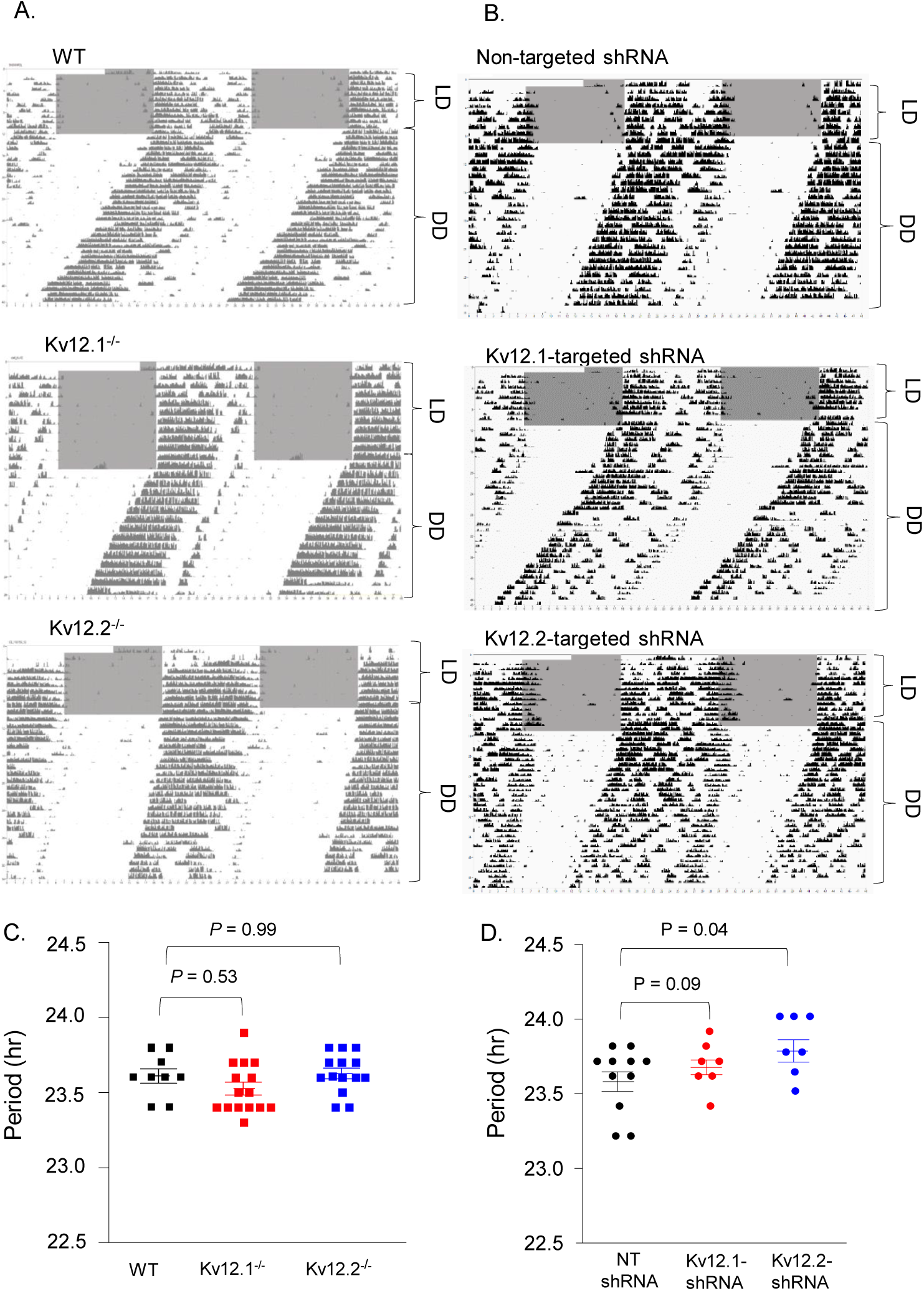
Rhythmic wheel-running activity is maintained in Kv12.1^−/−^ and Kv12.2^−/−^ mice and with acute *in vivo* shRNA-mediated knockdown of Kv12.1 or Kv12.2 in adult animals. Representative recordings of wheel-running activity of WT (top), Kv12.1^−/−^ (middle) and Kv12.2^−/−^ (bottom) mice (A) and WT mice that received bilateral SCN injections of the non-targeted (top), Kv12.1-targeted (middle), or Kv12.2-targeted (bottom) shRNA-expressing AAV8 (B). Continuous recordings were obtained for ∼10 days in 12:12 hr LD (indicated by the grey and white backgrounds, respectively) conditions, followed by at least 20 days in DD (constant darkness, indicated by the white background). The circadian periods, measured in each WT (▪), Kv12.1^−/−^ (▪), and Kv12.2^−/−^ (▪) mouse and in each animal expressing the NT (●), Kv12.1-targeted (●) and Kv12.2-targeted (●) shRNA in DD, are plotted; P values (one way ANOVA are indicated). (C) The mean ± SEM circadian periods of locomotor activity determined in WT (n = 9), Kv12.1^−/−^ (n = 15) and Kv12.2^−/−^ (n = 14) are similar. The mean ± SEM circadian periods (D) of locomotor activity measured in animals expressing the Kv12.1-targeted shRNA (n = 7), the Kv12.2-targeted shRNA (n = 7) and the non-targeted shRNA (n = 11) are also similar.

## Discussion

The results of the experiments detailed here demonstrate that the targeted deletion of Kv12.1 or Kv12.2 or the shRNA-mediated knockdown of Kv12.1 or Kv12.2 selectively increased the repetitive firing rates of SCN neurons at night, and eliminated the day-night difference in mean repetitive firing rates that is characteristic of WT SCN neurons.^[7–10]^ Direct application of the Kv12 channel blocker CX4^[30]^ increased repetitive firing rates at night, but not during the day, and the amplitudes of CX4-sensitive currents in WT SCN neurons were found to be higher at night than during the day. Additional experiments revealed that dynamic clamp-mediated subtraction of modeled I_Kv12_ selectively increased the repetitive firing rates of nighttime, and not daytime, SCN neurons. Despite the loss of day-night oscillations in mean repetitive firing rates, mice lacking Kv12.1 or/and Kv12.2 currents display daily rhythms in locomotor (wheel running) activity.

### Kv12 channels selectively regulate the nighttime repetitive firing properties of SCN neurons

The targeted deletion of *Kcnh3* (Kv12.2) or *Kcnh8* (Kv12.1) selectively increased the rate of spontaneous action potential firing of SCN neurons at night; the daytime repetitive firing rates of SCN neurons lacking Kv12.1/Kv12.2 are indistinguishable from WT cells. The higher repetitive firing rates in nighttime Kv12.1^−/−^ and Kv12.2^−/−^, compared with WT, SCN neurons are accompanied by higher input resistances and depolarized membrane potentials. The selective effects on nighttime firing rates and membrane properties were also observed in DKO (Kv12.1^−/−^/Kv12.2^−/−^) SCN neurons (Table 1). Acute, *in vivo* Kv12.1-targeted or Kv12.2-targeted shRNA-mediated knockdown of Kv12.1 or Kv12.2 in adult SCN neurons also selectively increased nighttime repetitive firing rates (Table 2). In addition to demonstrating a critical physiological role for Kv12-encoded K^+^ channels in regulating the nighttime firing and membrane properties of mature SCN neurons, these results also suggest that the firing properties of Kv12.1^−/−^, Kv12.2^−/−^ and Kv12.1^−/−^/Kv12.2^−/−^ SCN neurons do not reflect effects secondary to the loss of Kv12.1/Kv12.2 channels throughout development. Mimicking the effects of the Kv12.1/Kv12.2 knockout/knockdown, dynamic clamp-mediated subtraction of I_Kv12_ selectively increased the repetitive firing rates of WT nighttime SCN neurons. These results contrast with the effects of manipulating another subthreshold K^+^ current, I_A_: dynamic clamp-mediated subtraction of I_A_ increased spontaneous repetitive firing rates in WT nighttime *and* daytime SCN neurons.

### Relationship to previous studies

Several previous studies have demonstrated selective roles for other (i.e., non-Kv12-encoded) types of K^+^ channels in regulating the daytime and/or nighttime repetitive firing rates of SCN neurons. Acute *in vivo* shRNA-mediated knockdown of *Kcnd1*, which encodes the Kv4.1 α subunit and contributes to the generation of I_A_, in adult SCN neurons, affects daytime *and* nighttime I_A_ densities and spontaneous firing rates.^[19]^ The densities of I_A_ were lower in Kv4.1-targeted shRNA-expressing, compared with WT, SCN neurons during the day *and* at night.^[19]^ Knockdown of Kv4.1 also increased repetitive firing rates during the day *and* at night.^[19]^ In contrast with the effects of the knockdown or deletion of Kv12 channels, the loss of Kv4.1-encoded I_A_ channels did not eliminate day-night oscillations in the repetitive firing rates of SCN neurons.^[19]^ Also in contrast with the results here, *in vivo* knockdown of Kv4.1-encoded I_A_ channels advanced the time of daily activity onset by ∼1.8 hr and shortened the periods of wheel-running activity in constant darkness by approximately 0.5 hr. ^[19]^

In contrast with the effects of manipulating I_A_,^[16, 17, 19]^ targeted deletion of *Kcnma1*, which encodes the pore-forming *α* subunit of large conductance voltage- and Ca^2+^-dependent K^+^ (BK) channels, results in increased repetitive firing rates selectively in nighttime SCN neurons,^[20]^ markedly reducing the day-night oscillation in mean repetitive firing rates characteristic of WT SCN.^[7–10]^ The circadian periods of locomotor activity and clock gene expression in *Kcnma1^−/−^* animals, however, were indistinguishable from WT animals, although the circadian amplitudes of wheel-running activity and other behavioral outputs were reduced in *Kcnma1^−/−^*, compared with WT, mice during the day *and* at night.^[20]^ Opposite cellular effects were seen with overexpression of *Kcnma1* in the SCN, i.e., repetiitve firing rates were reduced in daytime, but *not* in nighttime, SCN neurons.^[47]^ The overexpression of *Kcnma1* and the reduction in daytime repetitive firing rates, however, was not accompanied by changes in the periods or the amplitudes of circadian behaviors.^[47]^

Previous studies have also demonstrated marked day-night differences in fast delayed rectifier (FDR) K^+^ current densities and in the expression of the *Kcnc1* and *Kcnc2* transcripts, which encode the Kv3.1b and Kv3.2 α subunits and generate FDR K^+^ currents, in SCN neurons.^[14]^ In addition, it was reported that exposure to the K^+^ channel blocker, 4-aminopyridine (4-AP) at a concentration of 0.5 mM, selectively reduces daytime repetitive firing rates and eliminates daily rhythms in the firing rates of SCN neurons.^[14]^ Similar effects on daytime firing rates were seen in SCN neurons from (*Kcnc1^−/−^*/*Kcnc2^−/−^*) animals lacking both Kv3.1b and Kv3.2,^[15]^ consistent with the suggestion that 0.5 mM 4-AP selectively affects FDR K^+^ currents in SCN neurons.^[14]^ Despite elimination of day-night oscillations in repetitive firing rates, daily rhythms in *Per2* expression were intact in *Kcnc1^−/−^*/*Kcnc2^−/−^* animals.^[15]^

### Molecular determinants of day-night differences in Kv12-encoded K^+^ current densities in the SCN

Consistent with the selective effects of the deletion or the acute knockdown of Kv12 subunits, action potential- (voltage-) clamp experiments revealed that the mean density of Kv12-encoded K^+^ currents in WT SCN neurons is much higher at night than during the day. The quantitative RT-PCR analysis completed, however, indicates that the day-night difference in Kv12-encoded K^+^ current densities does not result from circadian regulation of the *Kcnh3*/*Kcnh8* transcripts encoding the Kv12.1/Kv12.2 α subunits. Although it is certainly possible that Kv12.1/Kv12.2 protein expression in the SCN is rhythmic despite no day-night differences in *Kcnh3/Kcnh8* transcript expression, exploring this possibility must await the availability of validated anti-Kv12.1 and anti-Kv12.2 antibodies. It is also possible that Kv channel accessory subunits play a critical role in regulating day-night differences in Kv12-encoded K^+^ currents densities, as has been demonstrated for the accessory β2 subunit of *Kcnma1*-encoded BK currents.^[22]^

A rather large number of intracellular and transmembrane accessory subunits of Kv channels have been identified,^[55]^ and members of at least one subfamily, the *Kcne* subfamily, have been shown to regulate the expression and properties of Kv12.2-encoded K^+^ channels.^[48]^ Reducing *Kcne1* or *Kcne3* expression in *Xenopus* oocytes, for example, increased the amplitudes of heterologously expressed Kv12.2 currents, whereas over-expression of *Kcne1* or *Kcne3* reduced Kv12.2 current amplitudes.^[48]^ Additional experiments revealed that manipulating *Kcne1* or *Kcne3* affected the surface expression of Kv12.2, whereas total Kv12.2 protein was not affected.^[48]^ Although *Kcne1* and *Kcne3*, as well as other *Kcne* subfamily members, are expressed in the SCN (http://circadb.hogeneschlab.org), it is not clear whether any of these transcipts and/or the proteins encoded by these transcripts display daily rhythms in expression. It will be important to explore this possiblity directly, as well as to determine the functional consequences of manipulating *Kcne1* or *Kcne3* expression on the repetitive firing properties of SCN neurons.

Post-translational modifications of the Kv12.1 and/or Kv12.2 α subunits, or of accessory subunits of Kv12-encoded channels, could also play a role in modulating Kv12 current densities in SCN neurons by influencing cell surface channel expression directly or by modifying the conductance and/or the open probability of single Kv12 channels. In this context, it is of interest to note that *in vivo* constitutive activation of the two isoforms of the serine/threonine kinase glycogen synthase kinase 3, GSK3, previously shown to display rhythms in expression and phosphorylation of components of the molecular clock,^[49, 50]^ altered the repetitive firing rates of SCN neurons and disrupted rhythmic locomotor activity.^[51]^ Subsequent work revealed that GSK3 regulates the amplitude of the persistent component of the Na^+^ current, I_NaP_,^[52]^ in SCN neurons, and that GSK3 inhibition selectively reduced the spontaneous repetitive firing rates of SCN neurons during the day.^[53]^ It will be of interest to determine if GSK3, or downstream targets of GSK3, plays a role in regulating day-night differences in Kv12-encoded K^+^ current densities in the SCN.

### Physiological implications

The direct link between the spontaneous firing of action potentials and the output of the SCN was established with the *in vivo* demonstration that continuous infusion of the voltage-gated Na^+^ channel toxin, tetrodotoxin (TTX), into the SCN of unanesthetized and unrestrained animals (rats) disrupted the circadian rhythm of drinking.^[54]^ Additional *in vivo* experiments revealed a negative correlation between the repetitive firing rates of SCN neurons and wheel running activity, i.e., wheel running activity was high at night when repetitive firing rates were low and, conversely, wheel running activity was low during the day when repetitive firing rates were high.^[55]^ In addition, the acute silencing of SCN neurons during the day (by application of TTX) triggered wheel running, an observation interpreted as demonstrating that repetitive firing rates directly determine the temporal profile of behavioral activity.^[55]^ *In vitro* experiments on SCN explants also showed that application of TTX eliminated firing and circadian rhythms in vasopressin release.^[56]^

Given these combined observations, the finding here that the loss of day-night differences in the repetitive firing rates of SCN neurons with the loss/knockdown of Kv12.1/Kv12.2 did not affect the period of wheel-running activity seems surprising. As noted above, however, previous studies revealed that manipulating *Kcnc*-encoded FDR^[14, 15]^ or *Kcnma1*-encoded BK^[20, 47]^ K^+^ currents reduced/eliminated day-night differences in repetitive firing rates in the SCN, while having little or no effect on rhythmic wheel running activity.^[14, 15, 20, 47]^ One interpretation of the marked difference in the functional consequences of applying TTX versus changing individual K^+^ conductances is that the cellular effects are very different. Manipulating individual K^+^ currents, for example, may alter the daytime and/or nighttime repetitive firing rates of SCN neurons, but, unlike TTX, does not eliminate repetitive firing altogether. In addition, whereas application of TTX silences all SCN neurons, the impact of manipulating individual K^+^ currents might be highly variable across cells, depending on the repertoire and densities of the K^+^ and non-K^+^ channels contributing to shaping individual action potentials and repetitive firing patterns in different cell types.^[33, 57, 58]^ Indeed, the experiments here revealed that the densities of the CX4-sensitive K^+^ currents were quite variable among nighttime SCN neurons. In addition, ∼30% of nighttime SCN neurons were unaffected by CX4, indicating that the day-night switch in the repetitive firing properties of this subpopulation(s) of neurons is regulated by distinct (non-Kv12-encoded) K^+^ conductance(s). Given the diversity of cell types in the SCN, distinguished based on morphological features, transmitter phenotypes, synaptic connectivities, and firing properties,^[2-4, 8-10, 33, 36, 57, 58]^ future studies aimed at identifying the critical ionic conductances regulating the intrinsic membrane properties and the day-night oscillations in the repetitive firing rates of SCN should be conducted on identified SCN cell types.

## Acknowledgements

The authors wish to thank colleagues in the Nerbonne and Herzog laboratories for helpful discussions and Mr. Richard Wilson for technical assistance. We also thank Dr. Randall Rasmusson of Cytocybernetics for assistance with the development of the I_Kv12_ and I_A_ models and the implementation of these models in dynamic clamp recordings from SCN neurons. Financial support provided by the National Institute of General Medical Sciences (R01 GM104991 to EDH and JMN) is also gratefully acknowledged. TOH was supported in part by a postdoctoral fellowship award from the United Negro College Fund-Merck Science Initiative. All interfering RNAs (RNAi) were obtained from the Washington University RNAi Core, supported by the McDonnell Genome Institute and the Children’s Discovery Institute (CDI-LI-2010-94), and all viruses were generated in the Hope Center Viral Vectors Core, supported by a Neuroscience Blueprint Core grant (P30 NS057105).

---

**Supplemental Figure 1.**
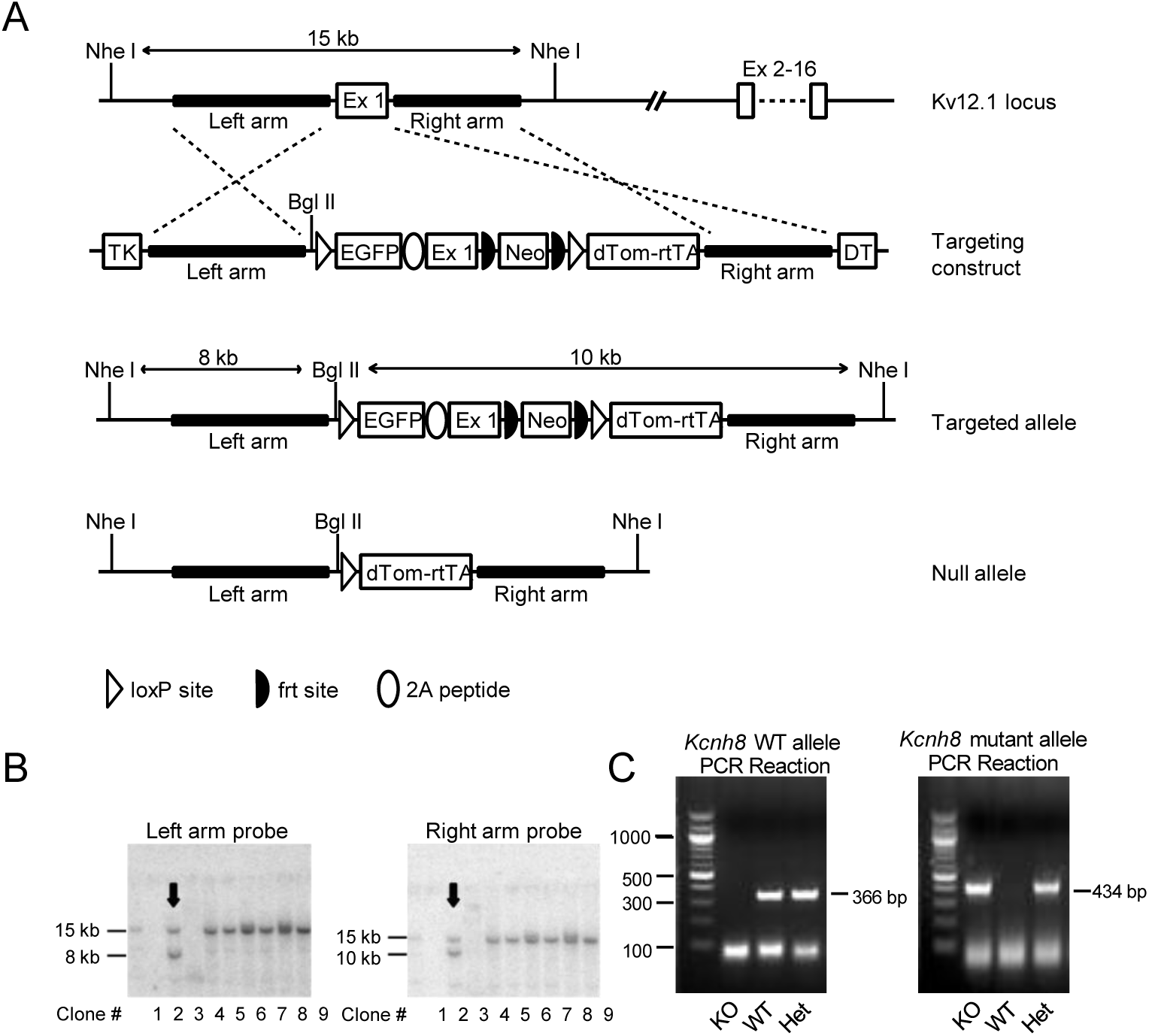
Generation of (Kv12.1^−/−^) mice harboring a targeted disruption of the *Kcnh3* locus. (A) Schematic of the targeted exon 1 *Kcnh8* (Ex1) locus, the linearized targeting construct, the initial targeted allele and the null (knockout) allele generated by Cre-loxP recombination. Targeting of the mouse *Kcnh8* (Kv12.1) locus involved homologous recombination (dashed lines) in mouse embryonic stem cells^53^ between the native *Kcnh8* locus and the targeting vector, and insertion of a loxP site and myristoyl-EGFP into Ex1 immediately upstream of the translation start. A viral 2A sequence joins the myristoyl-EGFP open reading frame to the Kv12.1 open reading frame to potentially allow EGFP-labeling of Kv12.1-expressing cells carrying the targeted allele. Downstream of Ex1 in the first intron, insertions include an frt-bracketed neomycin resistance cassette (Neo) driven by the PGK promoter for positive selection of targeted ES cells on G418, a 2^nd^ loxP site, and dTomato-2A-rtTA (reverse tetracycline trans-activator) cassette (dTOM-rtTA) which includes an SV40 polyadenylation sequence to terminate transcription and block expression of downstream exons. TK (thymidine kinase) and DT (Diptheria toxin) expression cassettes flank the left and right, respectively, arms in the targeting construct. These negative selection cassettes are eliminated by homologous recombination, but were included to suppress random insertion of the targeting construct into the ES cell genome. Note the native allele is bracketed by Nhe I restriction sites ∼ 15Kb apart. In the targeted allele, the Nhe I sites are preserved, but the distance between them is increased to ∼ 18 Kb, and a unique Bgl II site is introduced upstream of the 5’ loxP site. Hybridization probes located between either (left or right) arm andthe neighboring Nhe I site will label a ∼15 Kb band in a Southern blot of Nhe I/Bgl II digested genomic DNA for the WT allele. The same probes will label bands of 8 Kb (left arm) and 10 Kb (right arm) for the targeted allele. (B) Southern blot analysis of nine G418-resistant ES cell clones following Nhe I/Bgl II digestion of genomic DNA with a probe upstream of the left arm (left) or downstream of the right arm (right). The WT allele is identified by the 15 kb band with both probes, whereas the targeted allele is identified by 8 kb (left arm) and 10 kb (right arm) bands. Arrows indicate the ES cell clone positive for carrying the targeted allele used to generate the Kv12.1^−/−^ mouse line. Note DNA isolation failed for two clones (clones # 1 and 3). (C) ES clone #2 was karyotyped to confirm chromosome number and morphology and used for injection into C57BL/6J blastocysts. Three male chimeric mice were obtained, two of which transmitted the targeted allele through the germline. Mice carrying the targeted allele were bred with C57BL/6J-TgN(Zp3-Cre)93Knw females which express Cre-recombinase in the germline^54^ to generate heterozygous Kv12.1^+/-^ animals, which were then bred to generate Kv12.1^−/−^ mice. For genotyping, a two-step PCR reaction was used with primers specific to the WT (sense: TGGTCACAGTGCAGCGGCCAGGGAGTA and antisense: AAATTATTGCGCGGATGGAAAC AGAGGA) and targeted (sense: GTCACAGTGCAGCGGCCAGGGAGTAGC and aAntisense: CTTGGCGGTCTGGGTGCCCTCGTAGG) alleles for the *Kcnh8* gene. Bands at both 366 bp (WT) and 434 bp (targeted) identified heterozygous Kv12.1^+/-^ mice; bands at only 366bp or 434bp identified WT or homozygous Kv12.1^−/−^ mice, respectively.

**Supplemental Figure 2.**
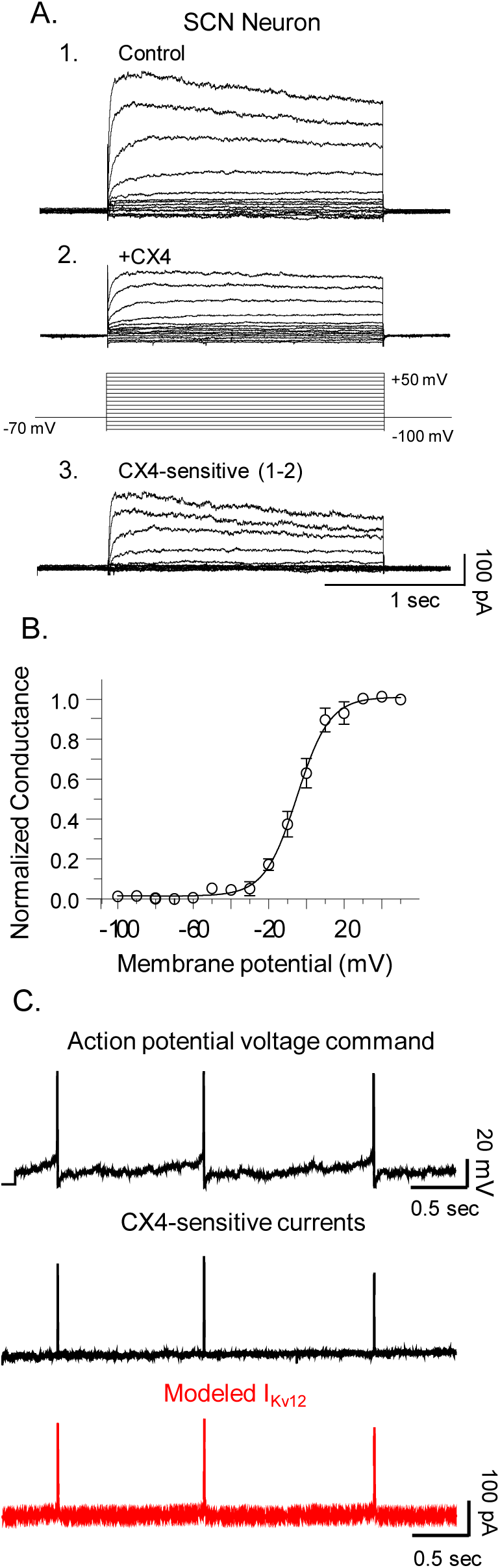
CX4-sensitive currents in WT SCN neurons and Cybercyte modeled I_Kv12_. (A) Representative whole-cell Kv current recordings obtained from a SCN neuron in an acute slice prepared from a night-phased (ZT19-ZT24) WT SCN are shown. Whole-cell Kv currents, evoked during (2s) voltage steps to potentials ranging from -100 to +50 mV (in 10 mV increments) from a holding potential of -70 mV, were first recorded in ACSF bath solution with 10 mM TEA and 10 mM 4-AP added (A1) and again following superfusion of the 10 mM TEA- and 10 mM 4- AP-containing ACSF with 20 uM CX4 added (A2). The voltage-clamp paradigm (in gray) is illustrated below the current records. Offline digital subtraction of the records obtained in the presence (A2), from the currents recorded in the absence (A1), of 20 μM CX4 provided the CX4-sensitive currents (A3). I_CX4_ conductances at each test potential were calculated and normalized to the maximal conductance (G_max_), determined in the same cell. (B) The mean ± SEM normalized conductances of activation of the CX4-sensitive currents are plotted as a function of the test potential and fit with single Boltzmanns. The V_1/2_ and the *k* values derived from these fits for current activation were V_1/2_ = -4.9 ± 1.0 mV; *k* = 8.7± 1.9 (n = 21). (C) These parameters were used to tune the I_K12_ model to fit the CX4-sensitive currents that were recorded during action potential-clamp experiments (see: Figure 4). The properties of currents produced by the Cybercyte I_Kv12_ model (lower panel, red) reliably reproduce the CX4-sensitive currents measured in action potential-clamp recordings (middle panel, black) from WT SCN neurons.

**Supplemental Figure 3.**
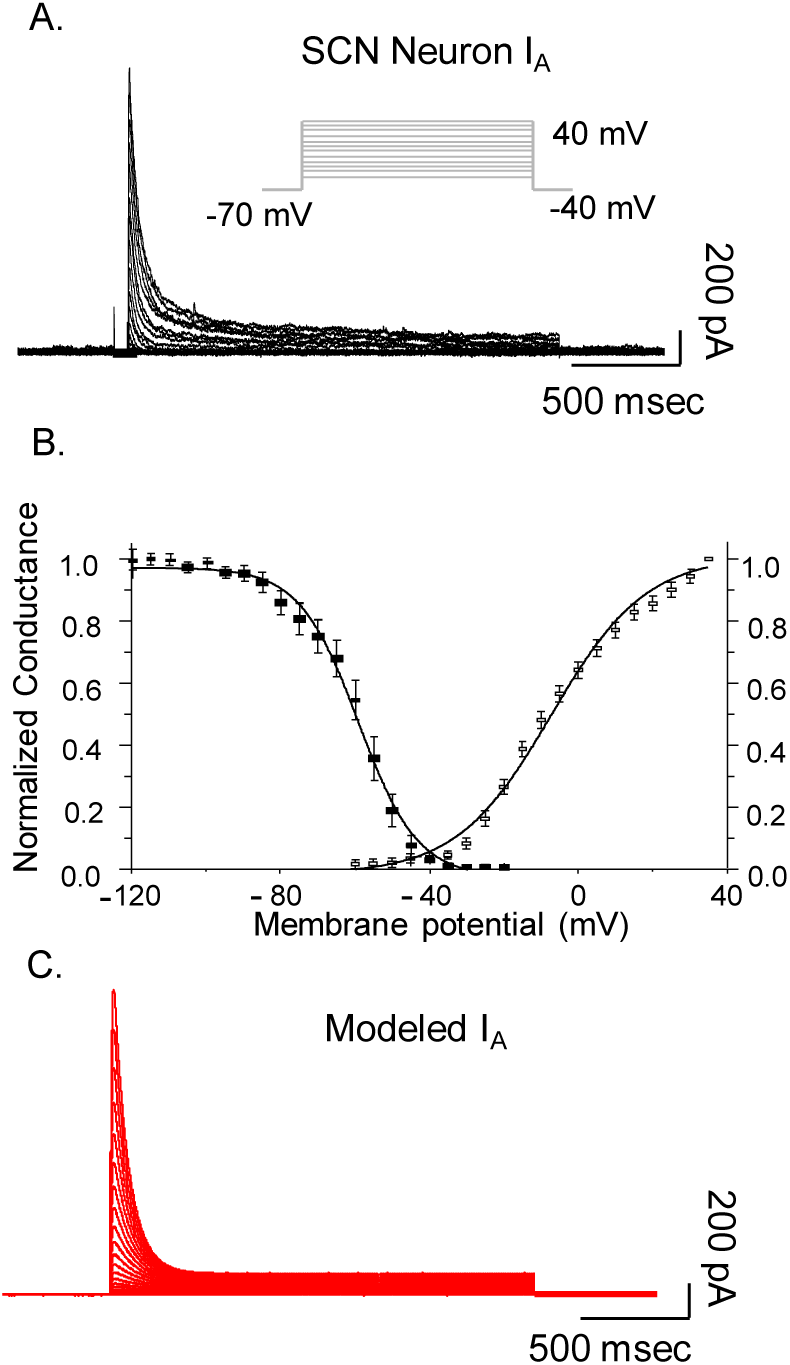
Cybercyte modeled I_A_. A Markov model describing the gating of the K^+^ channels that generate I_A_ in WT SCN neurons was developed based on a previously described model of the rapidly activating and inactivating, I_A_-like, K^+^ current in (ferret) ventricular myocytes,^[38]^ and was populated using previously acquired voltage-clamp data detailing the time- and voltage-dependent properties of I_A_ in mouse SCN neurons.^[19]^ (A) Representative I_A_ waveforms recorded from a WT SCN neuron^[19]^ in response to voltage steps to test potentials ranging from -40 mV to +40 mV (in 5 mV increments) from a HP of -70 mV are shown; the voltage-clamp paradigm is illustrated above the current records. The voltage-dependences of activation and inactivation for I_A_ in WT SCN neurons^[19]^ were determined using protocols identical to those described above for the CX4-senstive currents. The V_1/2_ and the *k* values derived from these fits for current activation (open symbols) and inactivation (closed symbols) were V_1/2_ = -9.3 ± 1.3 mV; *k* = 12.7 ± 0.8 (n = 12) and V_1/2_ = -59.4 ± 2.1 mV, *k* = 7.9 ± 0.7 (n = 12), respectively. (C) These parameters were used to tune the model. (C) The waveforms of the currents produced by the Cybercyte I_A_ model reliably reproduced I_A_ recorded from WT SCN neurons (A).

**Supplemental Figure 4.**
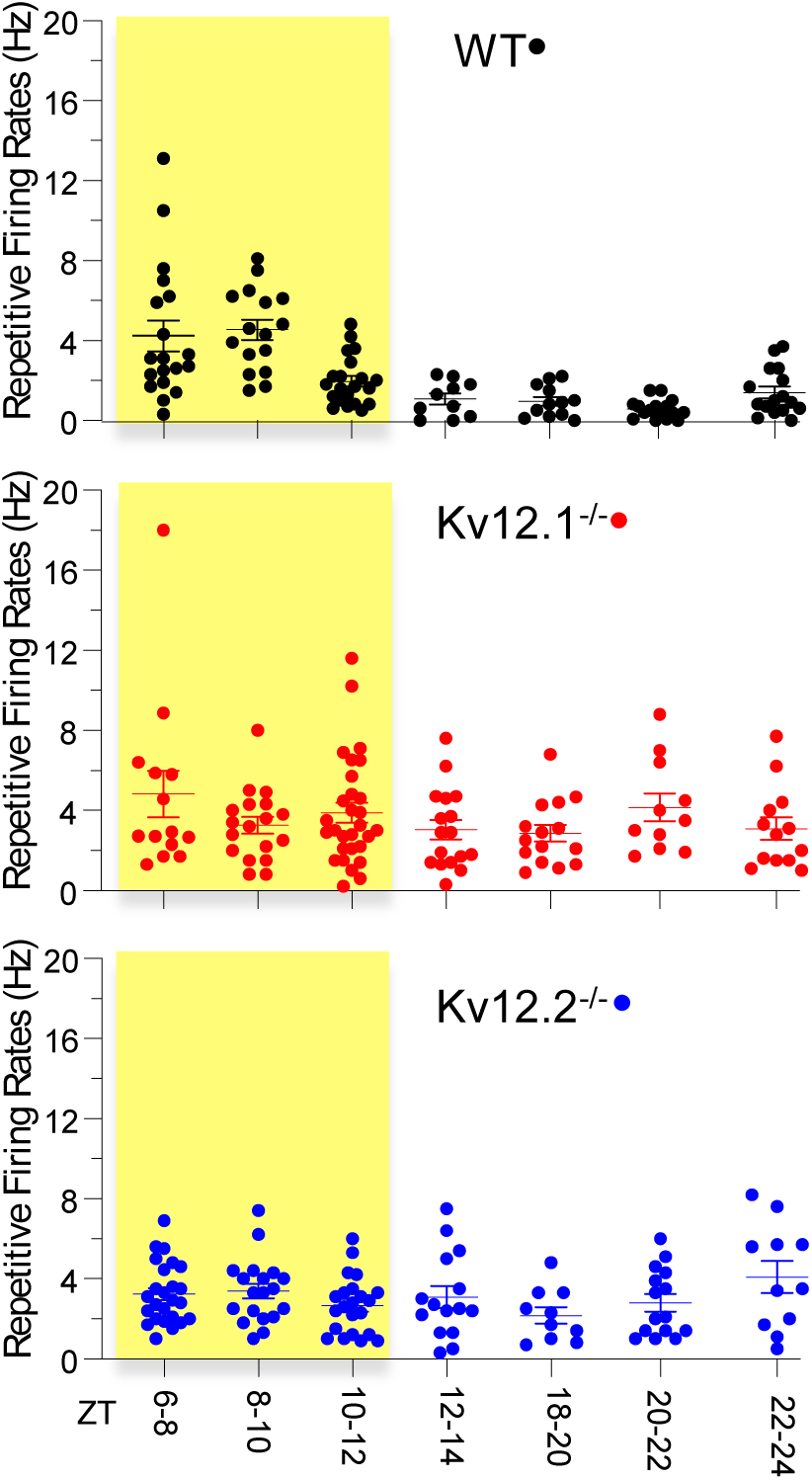
Repetitive firing rates of Kv12.1^−/−^ and Kv12.2^−/−^ SCN neurons are higher than WT SCN neurons throughout the night. Cell-attached recordings voltage recordings were obtained WT, Kv12.1^−/−^ and Kv12.2^−/−^ SCN neurons in acute slices prepared at various times throughout the circadian cycle. As is evident, mean ± SEM peak firing rates were high throughout the day (yellow shaded region) in WT (●) SCN neurons (n = 132), subsequently decreased during the transition from day to night, and remained low throughout the night. Conversely, the mean ± SEM peak firing rates of Kv12.1^−/−^ (●; n = 121) and Kv12.2^−/−^ (●; n = 125) SCN neurons did not vary measurably over time and were consistently high (similar to the daytime repetitive firing rates of WT SCN neurons) throughout the day and night.

**Supplemental Figure 5.**
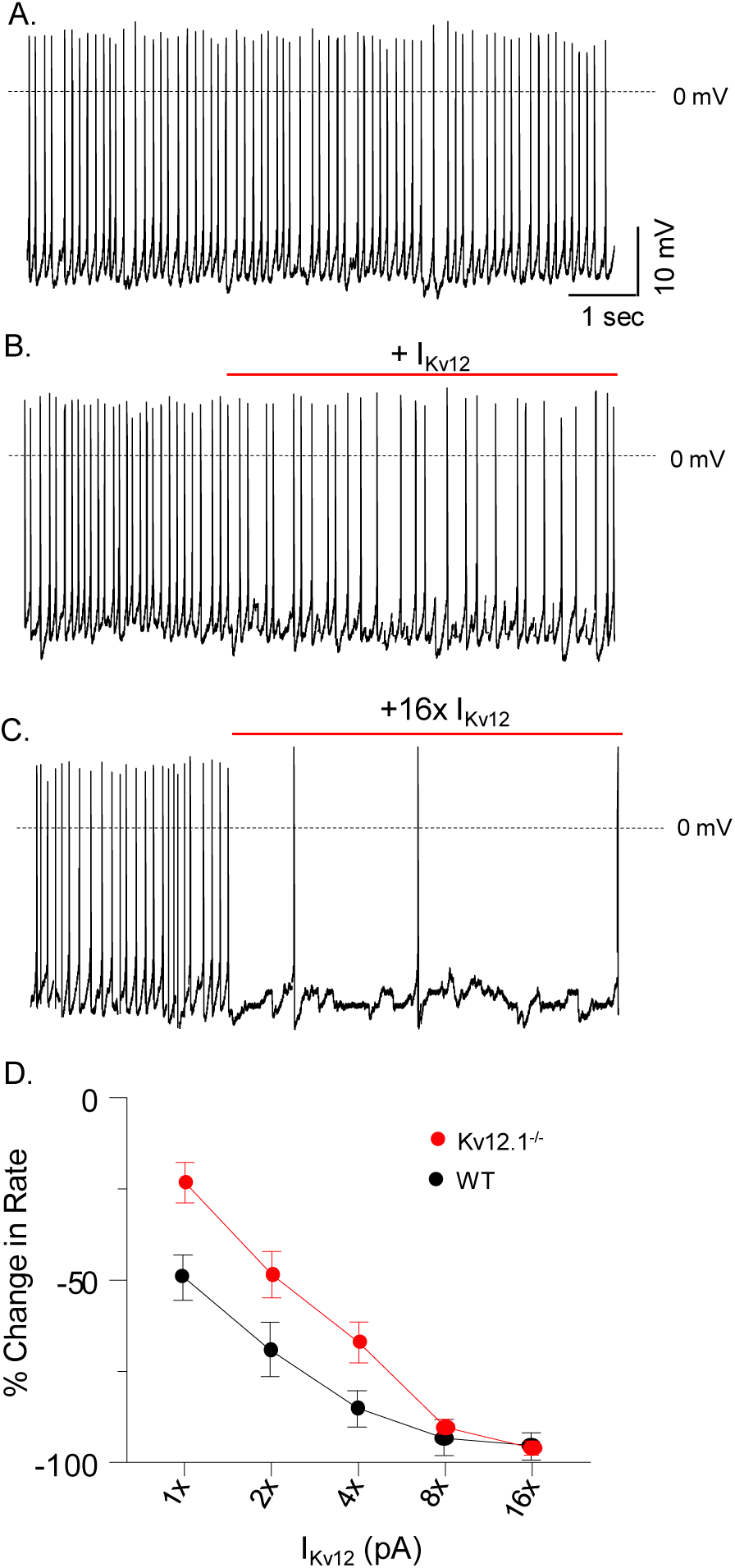
Dynamic clamp-mediated addition of I_Kv12_ decreases the rate of repetitive firing in nighttime Kv12.1^−/−^ SCN neurons. (A-C) Representative whole-cell current-clamp recordings obtained from a nighttime (ZT18-ZT20) Kv12.1^−/−^ SCN neuron under control conditions (A) and with dynamic clamp-mediated addition of modeled I_Kv12_ (B,C). Similar to WT SCN neurons (see Figure 7), the addition of modeled I_Kv12_ (B,C) reduced the repetitive firing rates of Kv12.1^−/−^ SCN neurons in direct proportion to the amplitude of the injected current. (D) The mean ± SEM (n = 11) percentage changes in the repetitive firing rates of nighttime Kv12.1^−/−^ SCN neurons (●) are plotted as a function of the amplitude of I_Kv12_ added. The percent changes in the repetitive firing rates of WT neurons with the addition of I_Kv12_ (from Figure 7D) are replotted here (●) to facilitate direct comparison of the results in WT and Kv12.1^−/−^ SCN neurons.

**Supplemental Figure 6.**
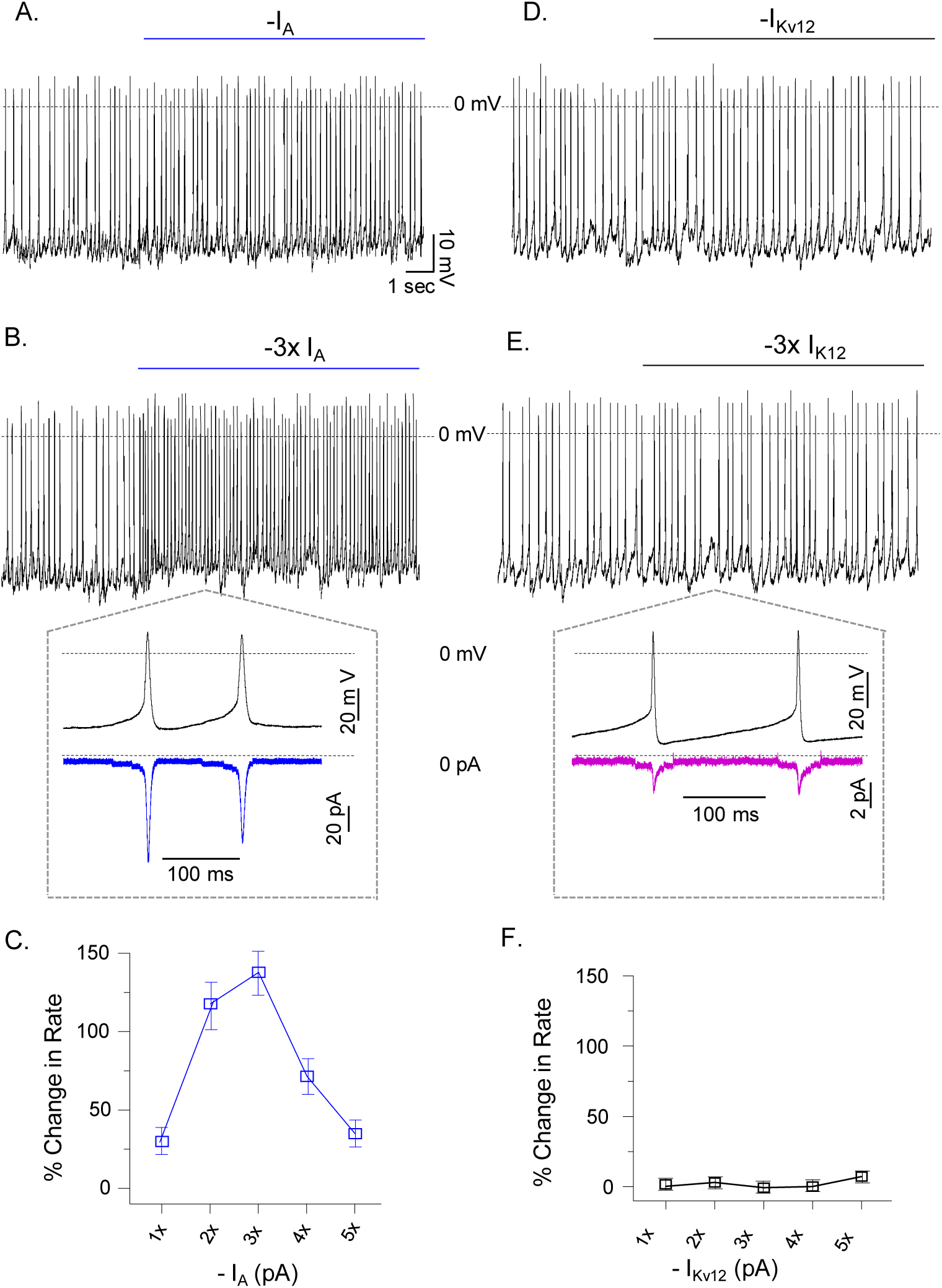
Direct comparison of the effects of dynamic clamp-mediated subtraction of I_Kv12_ versus I_A_ in daytime WT SCN neurons. (A,B,D,E) Representative whole-cell current-clamp recordings obtained from a day-phased (ZT7-ZT12) WT SCN neuron with dynamic clamp-mediated subtraction of modeled I_A_ (A,B) or I_Kv12_ (D,E). Subtracting modeled 1x or 3x I_A_ increased the rate of repetitive firing (A,B), whereas subtracting modeled 1x or 3x I_Kv12_ (D,E) in the same cell did not measurably alter firing rate. In the *insets* below panels B and E, the waveforms of individual action potentials (black), recorded in a representative WT daytime SCN neuron with subtracted modeled (-3x) I_A_ (B) or (-3x) I_Kv12_ (E), are plotted on an expanded timescale. The modeled (-3x) I_A_ (blue, B) and (-3x) I_Kv12_ (purple, E) waveforms in these cells are shown; the zero current levels are indicated by the dotted lines. Similar results were obtained in 6 additional cells. Plotting the mean ± SEM (n = 7) percentage changes in the repetitive firing rates of daytime WT SCN neurons reveals the mark difference in the effects of subtracting I_A_ (C) versus I_Kv12_ (F).

**Supplemental Figure 7.**
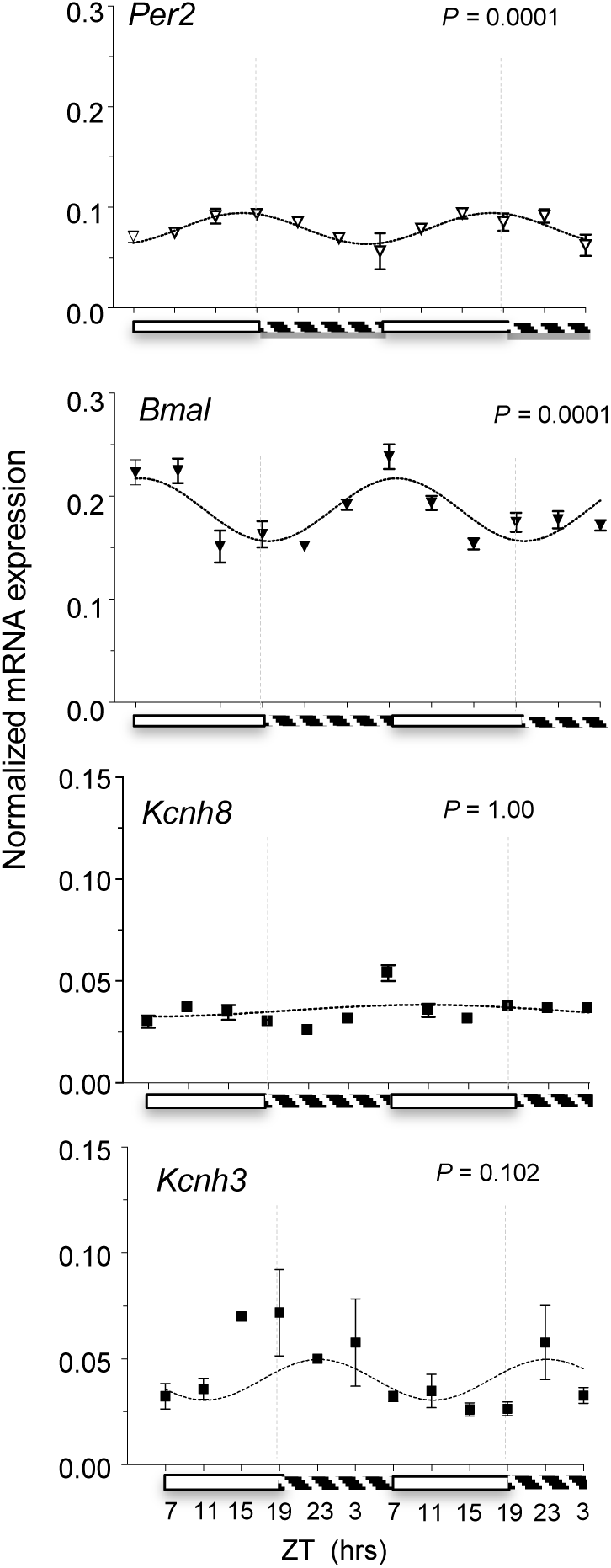
Expression of *Kcnh8* and *Kcnh3* in the SCN. The expression levels of the transcripts encoding *Per2*, *Bmal*, *Kcnh8* and *Kcnh3* were determined by quantitative RT-PCR analyses of SCN tissues samples, obtained every 4 hr over a 48 hr time period, from animals maintained in the standard and reversed 12:12 hr LD conditions, as described in **Materials and Methods**. The expression of each transcript was normalized to the expression of *Hprt* in the same sample. Mean ± SEM (n = 7 - 8) values for each transcript are plotted. Fitting the mean data for each transcript, using JTK cycle analysis^37^ with the period set to 24 hr, reveals that, as expected,^2,4,6^ the *Per2* (*P* = 0.0001) and *Bmal* (*P* = 0.0001) transcripts display 24 rhythms in expression. In contrast, the expression levels of the *Kcnh8* (*P =* 1.00) and *Kcnh3* (*P =* 0.102) transcripts do not display 24 hr rhythms (see text). The dashed lines on days 1 and 2 are at 7 pm.

